# Insight into spatial intratumoral genomic evolution in glioblastoma

**DOI:** 10.1101/2023.09.11.557112

**Authors:** Atul Anand, Jeanette Krogh Petersen, Lars van Brakel Andersen, Mark Burton, Martin Jakob Larsen, Christian Bonde Pedersen, Frantz Rom Poulsen, Peter Grupe, Torben A. Kruse, Mads Thomassen, Bjarne Winther Kristensen

## Abstract

Glioblastoma undergoes a complex and dynamic evolution involving genetic and epigenetic changes. Understanding the mechanisms underlying this evolution is vital for the development of efficient therapeutic strategies. Although treatment resistance is associated with intratumoral heterogeneity in glioblastoma, it remains uncertain whether hypometabolic and hypermetabolic lesions observed through positron emission tomography (PET) imaging are influenced by spatial intratumoral genomic evolution. In this study, we precisely isolated autologous hypometabolic and hypermetabolic lesions from glioblastoma using advanced neurosurgical and brain tumor imaging technologies, followed by comprehensive whole-genome exome and transcriptome analyses. Our findings revealed that hypermetabolic lesions evolved from hypometabolic lesions, harbored shrewd focal amplifications and deletions, and exhibited a higher frequency of critical genomic alterations linked to increased aggressiveness, upregulated APOBEC3 and hypoxic genes, and downregulated putative tumor suppressors. This study highlights spatial genomic evolution with diagnostic implications and unveils the obstacles and possibilities that should be considered in the development of novel therapeutic strategies.

**Statement of significance:** Glioblastoma is a multifaceted disease that is difficult to treat, and insights into the metabolic gradient observed in imaging and the underlying role of genomic evolution are lacking. This study is the first to investigate the molecular basis of hypermetabolic tumor lesions in glioblastoma using precise three-dimensional biopsy isolation, whole genome/exome, and mRNA sequencing. These findings have diagnostic significance, provide insights into therapeutic resistance, and shed light on the obstacles encountered by precision therapeutics for glioblastoma.

## Introduction

Glioblastoma (GBM) is the most common and aggressive brain tumor, with about 80,000 new cases per year worldwide and a median survival of about 15 months (1). Besides surgery, GBM treatment consists of radiation and chemotherapy, but most tumors inevitably recur (2). GBMs are characterized by extensive heterogeneity at both histological and molecular levels, representing the evolution of tumor aggressiveness and posing an obstacle to designing effective therapies (3–6). Although spatially random or bulk tumor biopsies have been used in analyses to understand the complexity of glioblastoma, they have contributed to our understanding of general GBM subtypes and their developmental trajectories to some extent (7–9). However, the link between intratumoral spatial metabolic heterogeneity and genomic evolution has not yet been fully explored.

In the clinical setting, magnetic resonance imaging (MRI) and positron emission tomography (PET) provide valuable information regarding tumor metabolism and location. However, it is unclear whether metabolically active tumor areas are genomically the most aggressive and whether this is associated with therapy resistance. We used MRI co-registered with ^11^C-methionine PET (^11^C-MET-PET) and ^18^F-fluorodeoxyglucose PET (^18^F-FDG-PET) brain imaging to identify tumor areas with different metabolic activities, followed by precise extraction of biopsies from these lesions using a neurosurgical stereotactic approach, guided by three-dimensional coordinates derived from MRI and PET imaging. The categorization of tumor areas in each patient was based on MRI, ^11^C-MET-PET, and ^18^F-FDG-PET. Those with low to medium PET intensity were classified as ‘‘hypometabolic lesions,” which is denoted by the Greek word ‘‘kryo” meaning cold, while those with high PET and MRI intensity were classified as ‘‘hypermetabolic lesions” which is represented by the Greek word ‘‘zesto” meaning hot. Finally, tissue areas with no intensity were labeled as ‘‘metabolically inactive” or ‘‘edge” in Greek, known as ‘‘akri”.

We embark on making a comprehensive analysis of genomic alterations in glioblastoma of kryo, zesto, and akri samples by performing exome and genome sequencing of tissue and blood samples to identify critical copy number variations, single nucleotide variants, gene fusions, somatic variants as well as RNA sequencing for functional analysis. By conducting these multi-omics analyses on multisectoral paired biopsies from the same patients with glioblastoma, allowing us to investigate intratumoral alterations, we identified a set of molecular alterations that may influence the regulation of glioblastoma biology and contribute to its heterogeneity and therapeutic resistance. This is the first study to perform a thorough genomic analysis on predetermined multi-regional metabolic tumor lesions from patients with glioblastoma, with an ideal experimental control setup.

## Results

### Clinicopathologic characteristics of metabolic stereotactic tumor lesions

We used a combination of MRI co-registered with ^11^C-MET-PET and ^18^F-FDG-PET to precisely geo-locate lesions. We employed a computer-guided stereotactic biopsy device to precisely obtain biopsies from the lesions. In total, we collected 23 biopsies, which consisted of six hypometabolic, ten hypermetabolic, one tumor satellite, and six peripheral lesions. Blood samples were obtained from six patients with glioblastoma (Figure 1A, Supplementary Figure S1 Table). Tumor regions with high-intensity of ^18^F-FDG and ^11^C-MET were classified as zesto, those with low uptake as kryo, and those with no uptake as akri (Figure 1B and Supplementary Figure S2). We obtained biopsies from the akri areas at the transition of the tumor to the normal brain parenchyma and from the non-eloquent (non-functioning brain tissue) areas outside the tumor.

**Figure 1.**
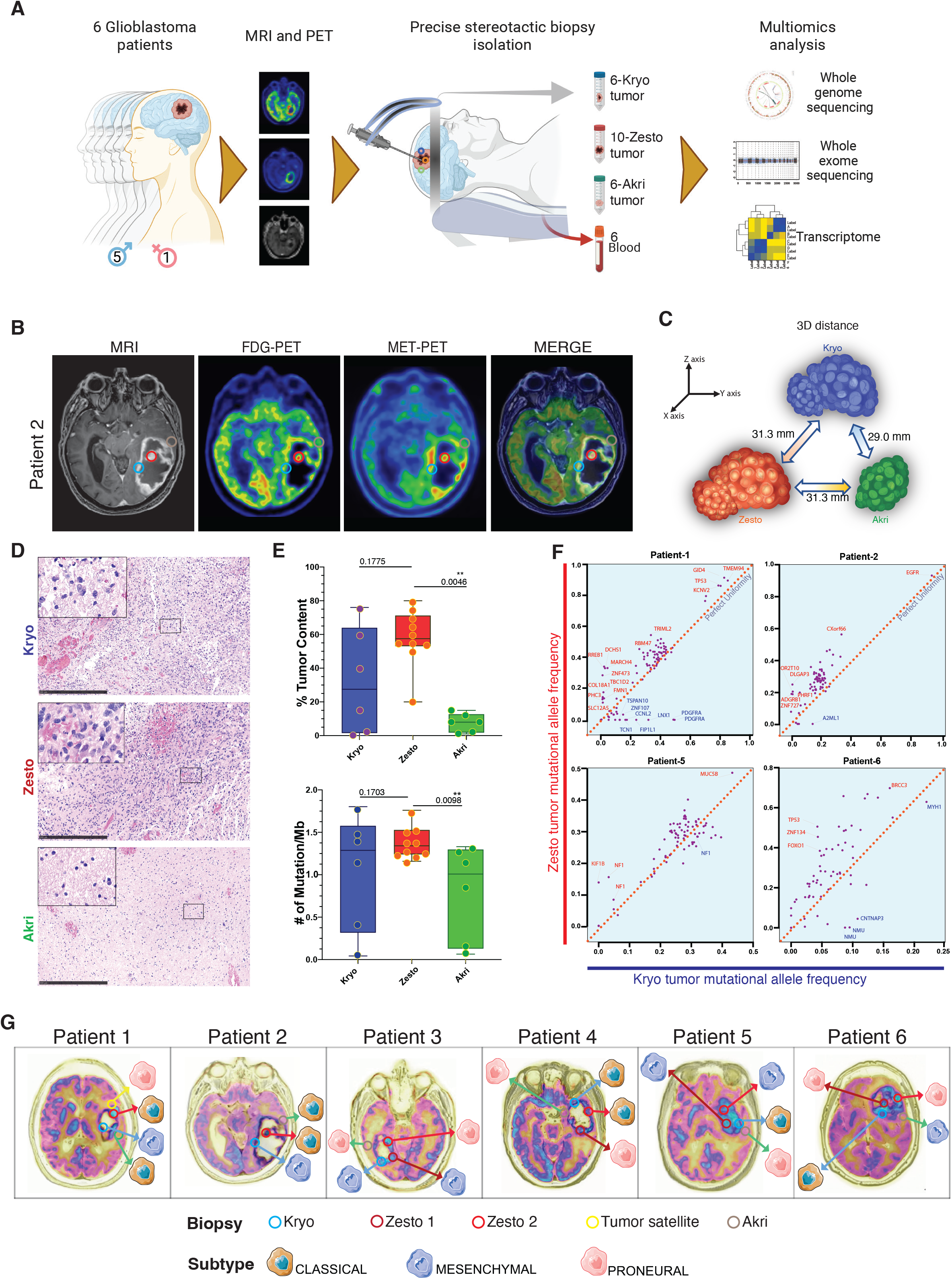
Summary of the experimental design and characterization of glioblastoma zesto, kryo, and akri tumor lesions: **(A)** Schematic representation of experimental setup to study molecular differences between hypometabolic (kryo), hypermetabolic (zesto), and no metabolic (akri) activity in glioblastoma lesions. **(B)** MRI, [^18^F] FDG-PET, [^11^C] MET-PET, and co-registration images from a representative glioblastoma patient (patient 3) showing the location of the kryo (blue circle), zesto (red circle), and akri (green circle) lesions. Blue represents the lowest, and red represents the highest accumulation of radiotracers. **(C)** Illustration representing the 3-dimensional distances between kryo, zesto, and akri lesions (other patient data are represented in supplementary figure S2). **(D)** Visualization of cellularity by HE-stained kryo, zesto, and akri lesions, scale bar 500µm. The inserts show the higher magnification of a representative area. **(E)** Box plots show exome analysis-based tumor fraction estimates (top) and analysis of tumor mutational burden in kryo, zesto, and akri lesions (bottom). Error bar limits are minimum to maximum; the center line is the median. An unpaired t-test determined statistical significance. **(F)** Tumor mutational allele frequency analysis by whole-exome sequencing in autologous kryo (x-axis) and zesto (y-axis) from four glioblastoma patients. Kryo and zesto-specific variants are marked respectively with blue and red (other patients are shown in Supplementary Figure S1A). Data points represent mutations (200X mean coverage), kryo, and zesto the red dotted line represents perfect uniformity. **(G)** Glioblastoma subtype predictions based on transcriptome analysis of individual metabolic lesions.

The average 3-dimensional spatial distance between autologous lesions ranged from 29 mm for kryo to akri, 31.3 mm for kryo to zesto, and zesto to akri lesions, which were measured using the StealthStation Surgical Navigation System (Figure 1C and Supplementary Figure S1B). Histological analysis of the individual lesions showed similar cancer cellularity between zesto and kryo lesions, whereas akri lesions had lower cellularity (Figure 1D). DNA extracted from each lesion and blood sample was used for whole-exome sequencing (WES) and whole-genome sequencing (WGS), and RNA was used for transcriptome analysis. The median coverage for WES was 100-fold, while that for WGS was 25-fold, indicating that the target regions were covered by 100 or 25 reads, respectively. Overall, 438 single-nucleotide variants (SNV) in the WES data and 700 structural variants (SVs) and 882 copy number variation (CNV) events in the WGS data were identified (Supplementary Tables S1 and S2). Tumor content and mutational burden analyses based on WES data showed no significant differences between kryo and zesto lesions. However, akri lesions had significantly lower tumor content than zesto lesions. Notably, we observed similar mutational calls in akri and kryo lesions, despite the low tumor content in akri lesions (Figure 1E).

Next, we performed a tumor mutational allele frequency analysis between kryo and zesto lesions with similar tumor content from autologous glioblastoma patients. We observed many differences despite a common pattern between autologous kryo and zesto in lesions from four patients (Patients-1,2,5 and 6). While analyzing SNVs, it became evident that the mutational patterns in hypermetabolic lesions (zesto) were abundant in certain subclonal variations when compared to metabolically hypoactive (kryo) lesions. These observations suggested that hypermetabolic lesions in glioblastoma evolved by acquiring more aggressive clones with specific driver variants (Figure 1F).

Further mutational allele frequency analysis revealed that certain mutations had a higher frequency in hypermetabolic lesions than hypometabolic lesions in most patients. For example, in patient 1, *RREB1, DCHS1,* and *MARCH4* mutations had a mean Variant Allele Frequency (VAF) of 35% in zesto lesions, whereas *PDGFRA, LNX1, FIP1L1,* and *CCNL2* mutations had higher VAFs in kryo lesions. Patient 2 had zesto lesion-specific mutations in *DLGAP3, OR2T10, PHRF1, ADGRB1,* and *ZNF727*, whereas *A2ML1* was exclusive to Kryo. Patient 5 had three variants of the tumor suppressor gene *NF1* in zesto lesions, and patient 6 had *TP53*, *ZNF134*, and *FOXO1* mutations. Two of the zesto (patient-3 and 4) lesions showed an exceptionally high degree of mutational variants compared to kryo, possibly because the paired kryo lesions had a lower tumor content (Supplementary Figure S1A).

Furthermore, we used predefined gene signatures to understand the transcriptional heterogeneity and subclass of these intratumoral lesions (8). We noticed a distribution of transcriptional subtypes with 8 proneural, 8 classical, and 7 mesenchymal subtypes across the 23 different lesions, in line with previous findings (9) (Figure 1G). We found that for the 10 hypermetabolic (zesto) lesions, four were proneural, three were mesenchymal, and three were classical. In contrast, half of the hypometabolic tumor (kryo) lesions were classical, and the other half were mesenchymal. For the six akri lesions, three were proneural, two were classical, and one was mesenchymal. Overall, the multisectoral lesions obtained from the six patients display distinct subtypes at the transcriptional level. Thus, hypermetabolic tumor (Zesto) lesions appear to be spatially more diverse than hypometabolic tumor (Kryo) and periphery (akri) lesions at both the genomic and transcriptomic levels.

### Glioblastoma hypermetabolic lesions evolve from a hypometabolic lesions

To explore chromosomal abnormalities, we used somatic copy number variations (SCNVs) and B allele fraction (BAF) analyses of autologous kryo, zesto, and akri lesions. Analysis of patient-1 showed three copies of chromosome 7 (allele ABB) in kryo, but only two copies of chromosome 7 (allele BB) were present in the zesto lesion (Figure 2A). This indicates that the flow of evolution occurred from the hypometabolic to the hypermetabolic tumor, as the A allele was lost in the hypermetabolic state. Additionally, in the zesto biopsy, we observed a specific loss of chromosome 4 and micro amplification of the chromosomal region 4q12. Remarkably, we observed deletion of chromosomes 4, 13, 19, and 22 only in kryo biopsy, which suggests parallel and spatially localized evolution of the hypometabolic area of the tumor. Additionally, a 4q12 micro amplification was detected in the akri biopsy, which may be linked to the recurrence of the tumor and migratory tumor cells. Indeed, we observed a marked increase in BAF for chromosomes 7 and 10 in all hypermetabolic lesions, as well as elevated BAF for chromosomes 1, 2, 3, 4, 5, 6, 11, 12, 13, 14, 15, 16, and 18, which varied among patients (Supplementary Figure S11 and S12). To visualize these key spatial differences, we mapped the chromosomal abnormalities on the merged transverse MRI and concordant FDG-PET images, which showed intratumoral heterogeneity between different autologous lesions (Figure 2B and Supplementary Figure 3). Overall, these findings indicated that hypermetabolic tumors evolved from hypometabolic regions, as schematically illustrated in the tumor evolution model (Figure 2C).

**Figure 2.**
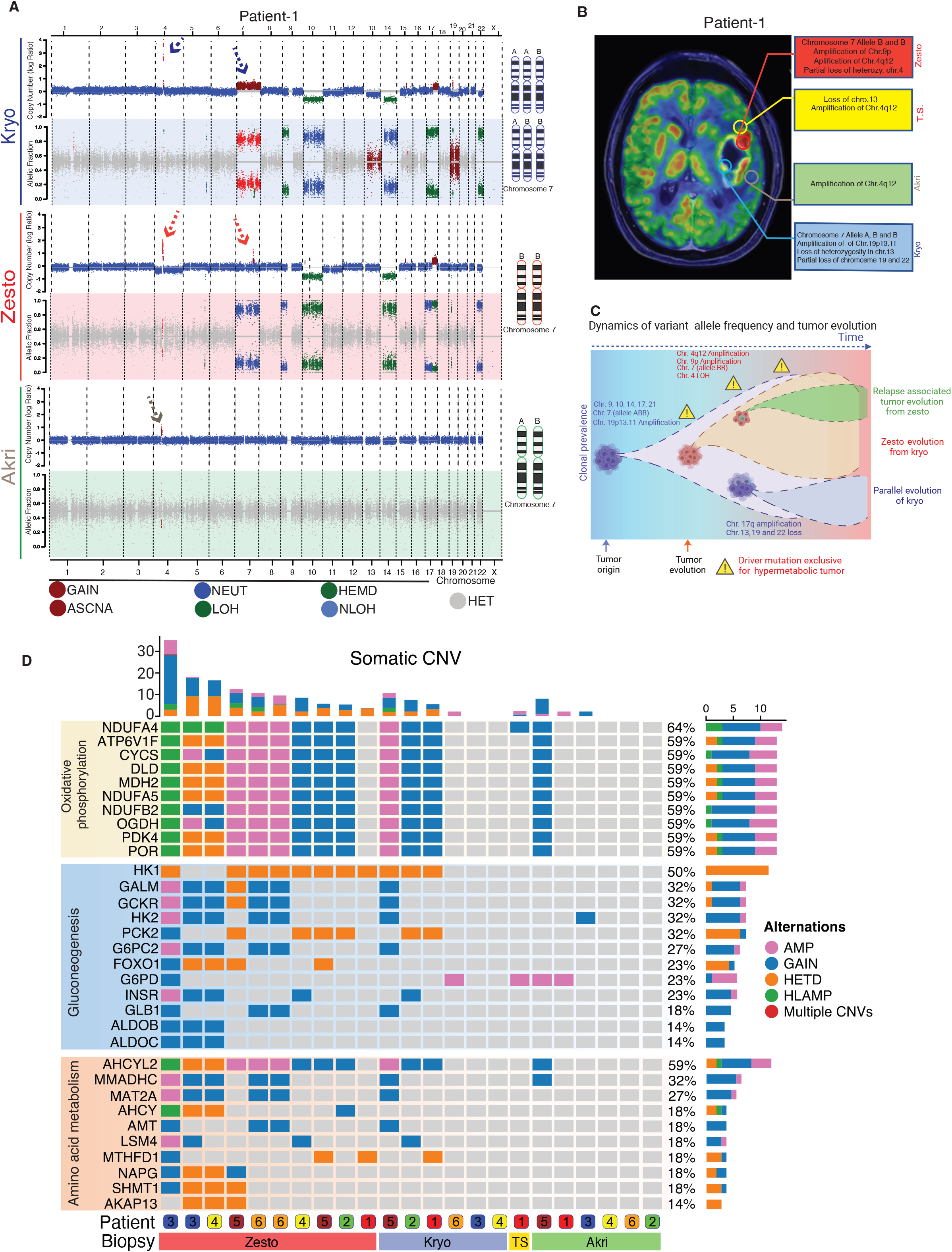
Hypermetabolic (Zesto) lesions evolve from hypometabolic (Kryo) lesions: Global copy number variation in autologous lesions from glioblastoma patient-1 as identified using the ASCAT (allele-specific copy number analysis of tumors) algorithm and tumor evolution model. **(A)** Copy number variation plot from whole genome sequencing (top) and B allele frequency (BAF) plot from whole exome sequencing analysis (bottom) from kryo, zesto, and akri lesions. The x-axis depicts the genomic location. Arrows on chromosomes 4 and 7 show focal amplification, and chromosomal aberrations, respectively. A schematic representation of the number of alleles from chromosome 7 is indicated on the right side of the plots. **(B)** Merged transverse MRI and FDG-PET images showing different geolocations of lesions. Textboxes represent specific chromosomal alterations in the autologous lesions from patient#1, and data from other patients is represented in supplementary figure S3. **(C)** Schematic tumor evolution model based on variant allele frequencies and chromosomal alterations illustrates the emergence of a zesto tumor from a kryo tumor and the parallel evolution of a kryo. Zesto lesion-associated focal amplification of chromosome-4q12 was also observed in the akri and tumor satellite, and these subclones of tumor in the akri lesion may contribute to tumor recurrence. Yellow triangles represent the driver mutations found only in the zesto biopsies. **(D)** Oncoplot showing the most prevalent putative driver mutations and CNAs in the metabolic zesto, kryo, and akri lesions of human glioblastoma grouped by metabolism pathways. (TS= tumor satellite).

Next we sought to outline specialized mutational developments through copy number alteration (CNA) analysis that may influence metabolic transformation to hypermetabolic tumors. Analysis of signature genes from oxidative phosphorylation, gluconeogenesis, and amino acid metabolism pathways revealed that 90% of zesto lesions had amplification, gain, and/or high-level amplifications of *NDUFA4, ATP6V1F, CYCS, DLD, MDH2* (oxidative phosphorylation), 40-60% of lesions had alterations of *GALM, GCKR, HK2, G6PC2, INSR* (gluconeogenesis) and 30-70% of lesions had alterations in *AHCYL2, MMADHC, MAT2A, AMT, LSM4* (amino acid metabolism) (Figure 2D). However, only 50% or less of kryo and akri lesions showed amplification or gain of genes associated with oxidative phosphorylation, gluconeogenesis, and amino acid metabolism. The presence of the most common mutations and CNAs in metabolic pathways in hypermetabolic lesions suggests that these mutations may have been acquired later in the tumor’s evolution to facilitate its higher metabolic demands, whereas a low frequency of these CNAs in some hypometabolic lesions may indicate their future transition to a hypermetabolic tumor.

### Hypermetabolic tumor lesions harbor shrewd focal amplifications and deletions

Next, we examined the heterogeneity of autologous lesions in patients with glioblastoma at the sub-chromosomal level. To achieve this, we analyzed chromosomal gains and losses in minimal common regions (MCRs) of chromosomes 4q12 and 19p13.11 using the Titan-CNA tool (10) (Figures 3A and 3E). Some of the substantial alterations identified by Titan-CNA in the 4q12 MCR covered recognized oncogenes and genes with known biological significance in glioma and glioblastoma, including *PDGFRA, REST,* and *CLOCK* (8,11,12) (Figure 3A dotted box). Interestingly, in a previous study, amplification of 4q12 was detected in 15% of GBMs cases (13). In addition, in the zesto biopsy, we detected amplified genes (*NMU*, *CHIC2*, *IGFBP7*, *TMEM165*, *PPAT*, *CEP135*, and *POLR2B*) that have not yet been studied in terms of its significance in glioblastoma (Figure 3B). Notably, genes such as *USP46, DCUN1D4, ERVMER34-1, THEGL, HOPX, SPINK2,* and *KIT,* which had a higher copy number in the akri and tumor satellite biopsy, can be associated with migration and recurrence of the tumor. Moreover, 19p13.11 MCR contained a region with a low copy number of *BABAM1, USE1, NWD1, DDA1, ANO8, OCEL1*, and *ABHD8*. To analyze the impact of these CNVs, we performed gene expression analysis of the 4q12 and 19p13.11 MCRs, which confirmed that these focal genomic alterations led to an altered mRNA expression for most of these genes, which can be seen in the chromosomal order in the respective lesions (Figure 3B and 3F). Indeed, genes with low copy numbers in the 19p13.11 MCR of zesto biopsy were found to have lower expression in glioblastoma tumors than in non-tumor samples (Figure 3G). These findings suggest that zesto tumors contain strategic, localized amplifications and deletions of genes that cause alterations in transcription, potentially affecting the metabolism and facilitating tumor evolution.

**Figure 3.**
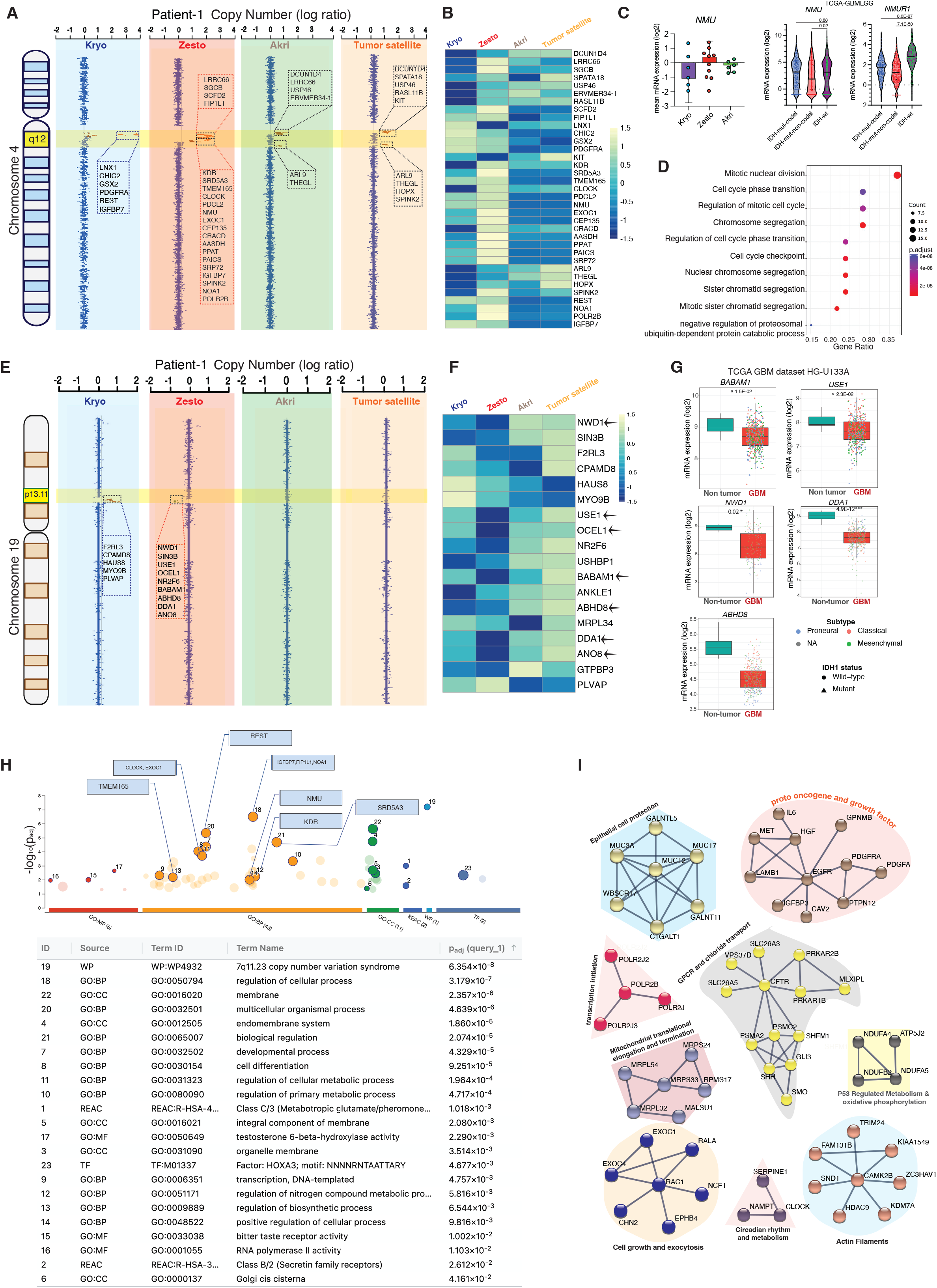
Focal amplification of genomic loci in kryo, zesto, akri, and tumor satellite lesions. **(A)** Copy number plots of chromosome 4q12 (log ratio) by whole-genome sequencing of the autologous kryo, zesto, tumor satellite, and akri lesions, dashed box shows the amplicons and list of involved genes. **(B)** mRNA expression of genes involved in amplicon found in zesto lesion. Genes are arranged in the chromosomal order to show the pattern of CNV and gene expression difference between autologous kryo, zesto, tumor satellite, and akri lesions. **(C)** Gene expression analysis of Neuromedin U (NMU) in the kryo, zesto, and akri lesions from different glioblastoma patients. Gene expression of NMU and Neuromedin U receptor 1 (NMUR1) from TCGA (The Cancer Genome Atlas) glioblastoma and low-grade glioma dataset. **(D)** Single gene ontology enrichment analysis of Neuromedin U for molecular function from glioblastoma and low-grade glioma dataset. **(E)** Copy number analysis of chromosome 19 by whole-genome sequencing of the autologous kryo, zesto, tumor satellite, and akri lesions. The yellow highlighted area shows an altered region in the Chr 19p13.11, and the dashed box shows the amplicons and a list of involved genes. **(F)** mRNA expression of genes involved in deletion and amplification found in Chr. 19p13.11 (patient-1). Genes are arranged in the chromosomal order to show the pattern of CNV and gene expression difference between autologous kryo, zesto, tumor satellite, and akri lesions. Zesto-specific deletions were marked with an arrow. **(G)** Gene expression of BABAM1, USE1, NWD1, DDA1, and ABHD8 from TCGA (The Cancer Genome Atlas) glioblastoma and non-tumor tissue samples. These genes are involved in zesto lesion-specific deletion in a glioblastoma patient-1. **(H)** G:Profiler-based gene set enrichment analysis performed on zesto lesion-specific CNVs from all the patients; some of the signature genes from the ontology terms are mentioned in the textbox. **(I)** A connectivity map of the significant protein-protein interaction involving genes from zesto lesion-specific CNVs from all six glioblastoma patients. 9 distinct functional clusters (illustrated in a discrete color) were detected, and disconnected nodes were excluded from the map.

### Neuromedin U as a putative hypermetabolic tumor driver

Neuromedin U (*NMU*) is a neuropeptide of the brain that is one of the genes amplified in the 4q12 MCR of glioblastoma hypermetabolic biopsy. *NMU* has recently emerged as an important neuropeptide that plays a key role in inflammation (14,15) and is underexplored in glioblastoma. We examined the mRNA expression of *NMU* in our cohort and observed a higher expression in hypermetabolic lesions. Moreover, *NMU* and its receptor *NMUR1* (neuromedin-U receptor 1) showed significantly higher mRNA expression in glioblastoma than in IDH-mutated 1p19q non-co-deleted samples (TCGA-GBM-LGG dataset) (Figure 3C). Single gene ontology enrichment analysis performed using Gliovis(16) (TCGA-GBM) on *NMU* revealed its possible role in cell division, cell cycle phase transition, and chromosomal segregation (Figure 3D). Moreover, single-cell sequencing analysis of *NMU* showed its expression in clusters of proliferative tumor cells; *MKI67*, *ZWINT*, *TOP2A*, *TK1*, and *CENPF*, which are predominantly expressed in proliferating tumor cells, were found to be co-expressed with *NMU* (Supplementary Figures 4A and 4B). Additionally, analysis of the IVYGAP dataset (http://gliovis.bioinfo.cnio.es/) indicated that NMUR1 was primarily expressed in the microvascular proliferation regions of the glioblastoma sections (Supplementary Figure S6F). This suggests the interdependency of tumor and vascular cells in glioblastoma hypermetabolic tumors, which may promote angiogenesis via Neuromedin U.

### The genomic landscape of hypermetabolic lesions predicts potential pathways for metabolism

Multiple minimal common regions (MCRs) have been identified in metabolic lesions (Supplementary Figures S7 A, B, C, D, and S8), which share similarities and differences among patients. Our global CNV analysis using VarScan (17) listed 1120 genes in the MCR regions of the genome from hypermetabolic lesions. To determine which pathways were enriched, we performed G:Profiler-based gene set enrichment analysis of biological processes and molecular functions. G:Profiler searches for a set of gene signifying pathways and GO terms (18). The most significant terms in the biological process category were regulation of cellular processes (FDR of 3.179×10^-7^), multicellular organismal processes, cell differentiation, and regulation of cellular and metabolic processes (Figure 3H). Additionally, the positive regulation of cellular processes, transcription, and nitrogen compound metabolism was significantly enriched in the zesto-biopsy-specific CNV gene set (Figure 3H). Subsequently, we performed protein-protein interaction (PPi) analysis using the STRING database to highlight the most significant pathways that may have roles in zesto lesions. With the minimum required interaction score of 0.900 (highest confidence), some of the PPi terms with the most significant values were transcription initiation, proto-oncogene, growth factor, G protein-coupled receptor (GPCRs), chloride transport, p53 regulated metabolism and oxidative phosphorylation, mitochondrial translational elongation and termination, and circadian rhythm (Figure 3I). Moreover, we evaluated hypermetabolic biopsy-specific CNVs (HS-CNVs) for copy number alterations and their impact on gene expression in the TCGA glioblastoma and low-grade glioma datasets. The samples with amplified genes and gain of chromosomes showed higher mRNA expression in the respective samples than in the diploid or homozygous deletion samples. (Supplementary Figures S4C). Strikingly, higher expression of HS-CNV signature genes was correlated with significantly shorter overall survival in patients with low-grade glioma and glioblastoma in the CGGA dataset (Supplementary Figures S4D). Our analysis revealed specific critical pathways and gene sets that may be essential for hypermetabolism and increased aggressiveness in tumor cells.

### Divergent somatic point mutations in the hypermetabolic lesions

We performed WES data-based single-nucleotide variant analysis using the Varscan2 variant caller, and subsequently used VarSeq to annotate, filter, and evaluate the identified variants (19). WES was conducted on six kryo, one tumor satellite, nine zesto, and six akri samples with autologous 6 blood samples to filter out passenger and background mutations from all six glioblastoma patients (Supplementary Table S3). To precisely evaluate somatic mutation frequency, distribution, and clonality, we used deeper read coverage for the exome, which gave us a high-confidence mutation calling and identified many significant somatic point mutations in different biopsy types (Supplementary Tables S2 and S4).

In our cohort, 438 somatic single-nucleotide variants (SNVs), in-frame insertions, and deletions were identified. The median exome-wide mutational burden was 1.4 mutations/Mb (Figure 1E), comparable to previous reports (20,21). Zesto lesions stand out as the most mutated samples, with no significant difference from kryo, but significantly higher than akri lesions. While performing an in-depth analysis of zesto lesions, we identified 21 genes with hypermetabolic-specific SNVs (HS-SNVs) (Figure 4A). Importantly, analysis conducted using the dbNSFP tool (database for non-synonymous SNPs functional predictions) (22) showed that multiple genes, including *COL18A1*, TBC1D2, PDIA2, *PIGO*, *OTULIN*, and *EMC7*, are predicted to induce damaging functional effects in hypermetabolic tumors. Conversely, certain variants (*CCDC38*, *OR5B3*, and *RTN1*) were predicted to have no discernible functional impact (Figure 4A and Supplementary Table S5).

**Figure 4.**
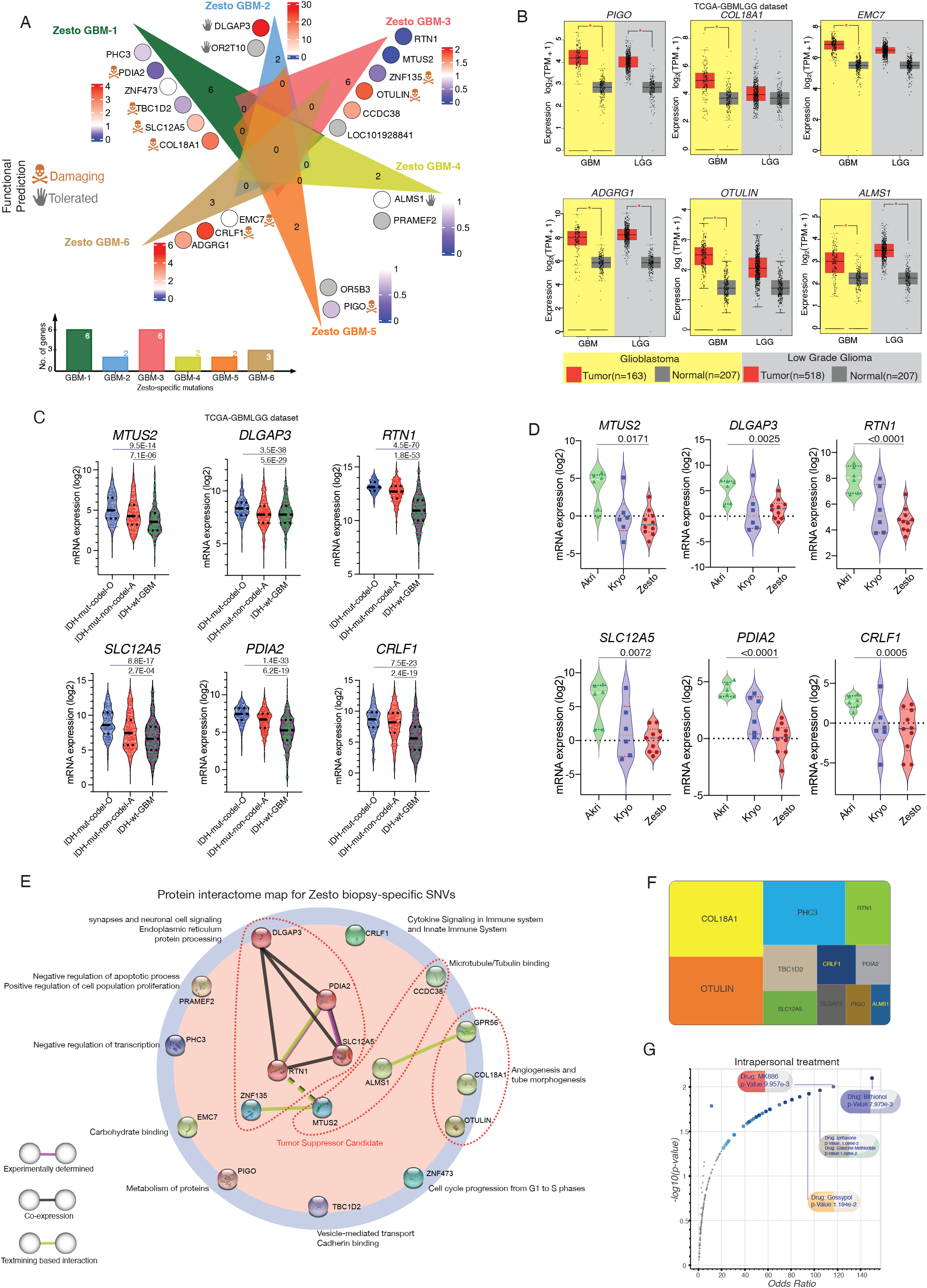
Single-nucleotide variant (SNV) profiles of hypermetabolic (zesto) lesions in glioblastoma patients. **(A)** Venn diagram representing the genes and number of somatic SNVs identified in different zesto lesions and its effect on mRNA expression from autologous patients with heatmap (color circles) showing fold change in the gene expression of the zesto compared to kryo lesions. The damaging functional effect of the SNVs is represented by the cranium sign. **(B)** mRNA expression of potential zesto lesion specific driver genes PIGO, COL18A1, EMC7, ADGRG1, OTULIN, and ALMS1 in low-grade glioma and glioblastoma from TCGA dataset. Statistical significance (P<0.01) was analyzed by one-way analysis of variance (ANOVA) and is indicated with an asterisk. **(C)** mRNA expression of metabolic zesto lesion-specific SNVs genes (*MTUS2, DLGAP3, RTN1, SLC12A5, PDIA2,* and *CRLF1*) in TCGA (The Cancer Genome Atlas) IDH-wildtype glioblastoma (GBM), IDH-mutated 1p19q co-deleted oligodendroglioma (O) and IDH-mutated 1p19q non co-deleted astrocytoma (A). These genes are significantly under-expressed in glioblastoma compared to lower-grade glioma. **(D)** mRNA expression of putative tumor suppressor candidate genes (MTUS2, DLGAP3, RTN1, SLC12A5, PDIA2, and CRLF1) found in WES analysis in the kryo, zesto, and akri lesions. Statistical differences in mRNA expression were assessed by a two-tailed t-test, the p-value indicates significance between akri and zesto lesions. **(E)** A connectivity map of the significant protein-protein interaction involving genes from zesto lesion-specific SNVs from all six glioblastoma patients. Three discrete functional clusters (illustrated with a dashed line) were detected, and disconnected nodes were placed on the circle with their potential biological role. **(F)** OncoScore plot showing the oncogenic potential of genes based on the text-mining algorithm by OncoScore tool, the bigger box size represents a higher OncoScore. A total of 8 genes (ZNF473, OR2T10, MTUS2, ZNF135, CCDC38, PRAMEF2, OR5B3, EMC7) had OncoScore of 0, indicating no prior association reported with cancer in the scientific journals, OncoScore ≥1, indicating at least one citation in a cancer-related study. **(G)** Volcano plot of drugs from the DSigDB gene set. Each point represents a single term, plotted by the corresponding odds ratio (x-position) and -log10(p-value) (y-position) from the enrichment results of the metabolic zesto lesion-specific gene set. Top 5 candidate drugs identified in drug-target enrichment using DSigDB gene datasets analyzed by Enrichr, the drug targets pathways contributed by zesto lesion-specific SNV genes. The larger and darker-colored the point, the more significantly enriched the input gene set is for the drug.

We used the NCI Genomic Data Commons (GDC) web tool to explore the presence and frequency of genes harboring HS-SNV in public datasets (23). The prevalence of mutations in the top 19 hypermetabolic biopsy-specific mutated genes was also analyzed in 57 cases (TCGA-GBM dataset), of which *ALMS1* (22%), *SLC12A5* (12%), *TBC1D2* (10%), *PRAMEF2* (8%), and *MTUS2* (8%) were the most common (Figure 5D). HS-SNV genes exhibit varying levels of genetic diversity and belong to different functional classes. To group them, we analyzed their mRNA expression in publicly available datasets for glioblastoma (GBM) and lower-grade gliomas (LGG) and divided them into two major categories. The first group, comprising *PIGO, COL18A1, EMC7, ADGRG1, OTULIN,* and *ALMS1*, demonstrated higher expression in GBM and lower-grade glioma than in normal brain tissue (Figure 4B). In contrast, the second group, consisting of *MTUS2, DLGAP3, RTN1, SLC12A5, PDIA2,* and *CRLF1*, exhibited significantly lower expression levels in IDH-wild-type glioblastomas compared to IDH-mutant 1p/19q-co-deleted oligodendrogliomas, IDH-mutant astrocytomas, and normal brain tissues. (Figure 4C and Supplementary Figures S5B). Importantly, transcriptome analysis data from this study showed that *MTUS2, DLGAP3, RTN1, SLC12A5, PDIA2*, *PHC3*, and *CRLF1* were significantly underexpressed in zesto lesions compared to akri lesions and had intermediate expression in kryo tumor lesions (Figure 4D). Moreover, single-cell mRNA expression analysis showed very low mRNA expression of *MTUS2*, *DLGAP3*, *PDIA2*, and *CRLF1* in the tumor cell cluster. Conversely, moderate expression levels were observed for *SLC12A5*, *RTN1*, and *PHC3* in both tumor clusters and oligodendrocyte progenitor cell (OPCs) clusters. (Supplementary Figure S5A).

**Figure 5.**
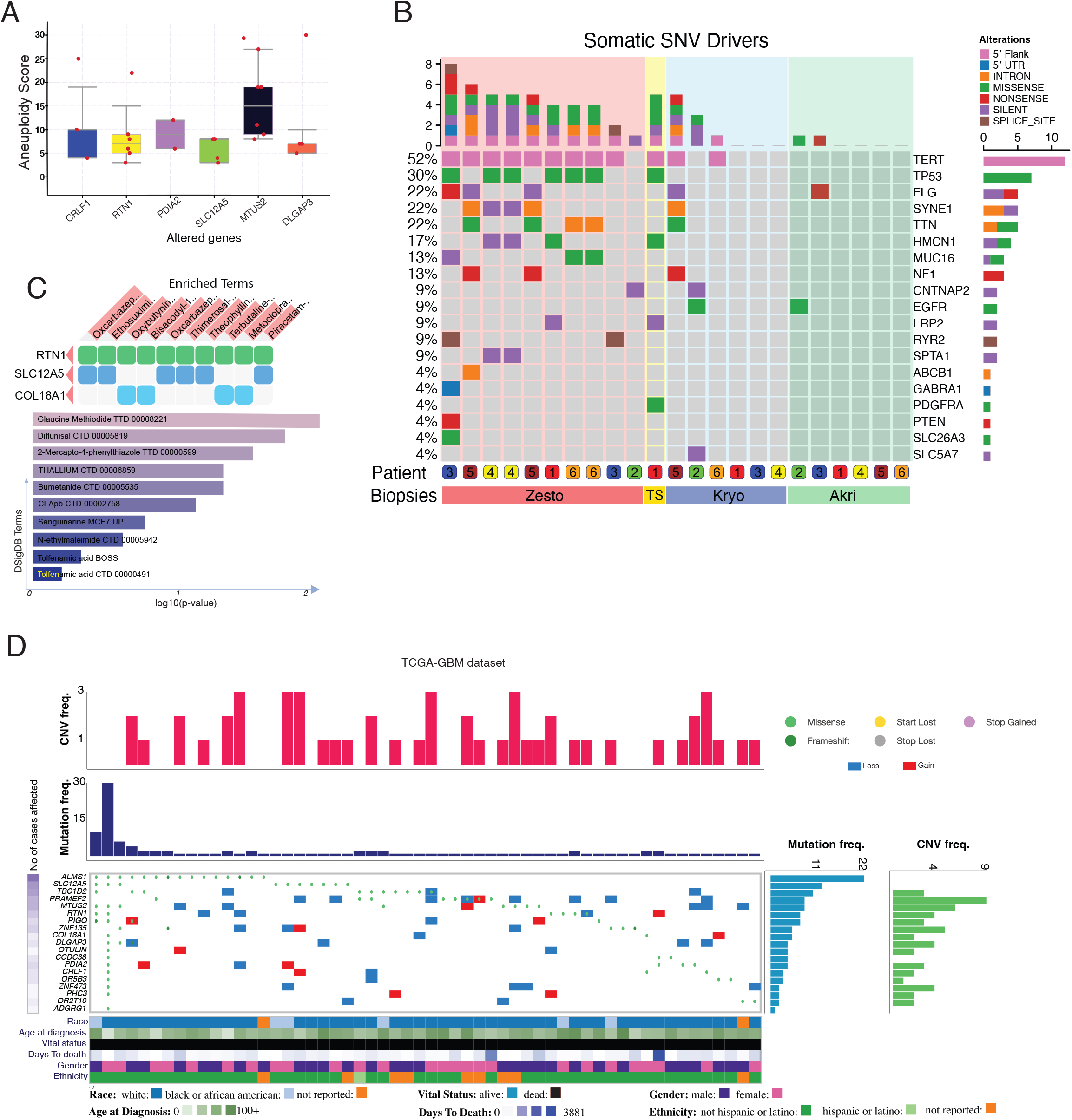
Candidate tumor suppressor genes with therapeutic and prognostic biomarker value. **(A)** Aneuploidy score of glioblastoma patients where zesto lesion-specific genes are altered, the y-axis indicates the aneuploidy score, and the x-axis indicates the altered genes, mutations in *MTUS2* show the highest aneuploidy level. Data was analyzed using the cbioportal tool. **(B)** OncoGrid plot from WES analysis showing genetic alterations (5′ flank, 5′ UTR, intron, missense, nonsense, silent, and splice site) of the most frequently mutated genes observed in glioblastoma. The top bar plot indicates the total no of genes altered in the corresponding patient lesion, and the right bar plot shows the SNV type and number of cases affected. The bottom color panel indicates the patient number and lesion type, TS represents the tumor satellite. **(C)** Heatmap and bar chart of top enriched drug and pathway terms from the DSigDB gene set library. The top 10 enriched most significant terms for the zesto lesion-specific gene set are displayed based on the -log10(p-value). The term at the top has the most significant overlap with the zesto lesion-specific gene set. **(D)** OncoGrid plot of genetic alterations (Missense, start loss, stop gained, frameshift, stop lost, and CNVs) from TCGA-GBM datasets analyzed on Genomic Data Commons (GDC) Data Portal, the plot shows the top 19 zesto lesion-specific genes and rare cancer drivers that have also been found mutated in 57 other GBM patients. The top bar plot (Red) indicates the frequency of CNVs in the corresponding patient tumor, 2nd the bar plot (navy blue) from the top shows the frequency of mutation in the related patient sample. The left heatmap represents the number of cases affected. Right sidebar plots indicate the overall frequency of mutation (blue) and CNV (green) in 57 glioblastoma patients. The grid in the center represents the type of mutation; red blocks indicate a gain of function, and blue indicates a loss of function. The bottom color panel indicates six classes of clinical information about the patients.

After observing a unique group of genes that may control the metabolism of glioblastoma in both our cohort and the public dataset, we evaluated their biological implications by conducting PPi analysis. STRING analysis of HS-SNV genes revealed the top three most prominent interacting pathways, including angiogenesis and tube morphogenesis (*GPR56, COL18A1*, and *OTULIN*), microtubule/tubulin binding (*MTUS2, ALMS1,* and *CCDC38*), and synapse and endoplasmic reticulum protein processing (*DLGAP3, SLC12A5, RTN1, PDIA2*, and *ZNF135*) (Figure 4E). Other terms involving genes were metabolism of proteins (*PIGO*), cytokine signaling in the immune system, innate immune system (*CRLF1*), carbohydrate binding (*EMC7*), and cell cycle progression from G1 to S phase (*ZNF473*). Our integrative analysis of HS-SNVs uncovered the potential functions of these genes, which could provide a better mechanistic understanding of hypermetabolic cancer cells.

To assess the oncogenic potential of HS-SNVs, we performed a text-mining algorithm-based OncoScore analysis (24), which indicated that *COL18A1, OTULIN, PHC3, TBC1D2, SLC12A5,* and *RTN1* were associated with an oncogenic potential to a high degree (Figure 4F), whereas *CRLF1, PDIA2, DLGAP3, PIGO,* and *ALMS1* were known to have an oncogenic role to a lesser degree. *ZNF473, OR2T10, MTUS2, ZNF135, CCDC38, PRAMEF2, OR5B3,* and *EMC7* had an OncoScore of 0, indicating no previous association with cancer in previous studies. These data highlight the potential role of HS-SNV genes in the regulation of angiogenesis, microtubule binding, synapse, and endoplasmic reticulum protein processing-related pathways.

### Hypermetabolic lesions as a contender for therapeutic intervention

To evaluate the selection of druggable targets in hypermetabolic tumors, we explored the therapeutic landscape across all HS-SNVs using the drug-target enrichment tool Enrichr, which employs the Drug SIGnatures Database (DSigDB) gene dataset (25). We found drug signatures of bithionol (p-value:7.973^-3^), MK886 (p-value:9.957e-3), and ipriflavone (p-value:1.095e-2) with the highest odds ratio (Figure 4G) predicted to target zesto-biopsy-specific SNVs genes. Furthermore, Enrichr analysis uncovered noteworthy druggability potential of oxcarbazepine, ethosuximide, and oxybutynin, as they demonstrated a substantial capacity to target reticulon 1, solute carrier family 12 member 5, and collagen type XVIII alpha 1 chain (Figure 5C). These findings suggest that bithionol, MK886 and oxcarbazepine possess the greatest potential for effective targeting of hypermetabolic lesions.

### Hypermetabolic lesion-specific SNVs are associated with genomic instability

As low-frequency cancer drivers and tumor suppressor-like genes play an essential role in tumor progression, we explored whether HS-SNVs have a cancer driver and tumor suppressor-like role, and how they correlate with glioblastoma genomic instability and patient prognosis. If HS-SNVs are driver genes, one of their effects may be an increase in chromosomal instability. An abnormal chromosome number or somatic DNA copy number is defined as aneuploidy which disturbs the vital cellular genomic balance. It is a characteristic of many aggressive tumors and is believed to initiate tumorigenesis (26). To study the effects of HS-SNVs on tumor suppressor-like effects in glioblastoma, we evaluated the aneuploidy levels of glioblastoma samples with mutations in HS-SNVs using the cBioPortal web tool (27,28). In sharp contrast, the mean aneuploidy score of MTUS2 was the highest (aneuploidy score of 15) among all HS-SNVs, whereas CRLF1, RTN1, and PDIA2 also had a moderate effect on aneuploidy levels (Figure 5A).

Based on the ability of HS-SNV’s suppress tumors, we speculated that the upregulation of these genes might have a negative effect on cell proliferation and metabolism. We performed correlation analysis to further validate the association between *MTUS2* and the glucose metabolism pathway. Correlations between *MTUS2* and *HK1, HK2, HK3, GALM, LDHA, GAPDH, ENO1,* and *PKM* expression were calculated. *MTUS2* was negatively correlated with most gluconeogenesis pathway genes, except *HK1* and *GAPDH* (coefficients > −0.20 and P<0.05) (Supplementary Figures S5C). Importantly, the protein expression levels of *MTUS2*, *SLC12A5*, and *RTN1* were strongly negatively correlated with the protein levels of MKI67 and Thymidine kinase 1(*TK1*), which are markers of cell proliferation and DNA replication. (all P<0.05, coefficients > −0.25 Supplementary Figures S5D). Moreover, we found that the mRNA expression of HS-SNVs genes (*MTUS2, RTN1, CRLF1, PDIA2,* and *DLGAP3*) was positively correlated in the glioblastoma cohort (Supplementary Figures S5E). This indicates that HS-SNVs genes (*MTUS2, RTN1, DLGAP3, CRLF1, PDIA2,* and *SLC12A5*) may play a role as putative negative regulators of gluconeogenesis and cell proliferation in hypermetabolic tumors.

Next, we aimed to elucidate whether the HS-SNVs signatures have prognostic value for the outcome of patients with glioblastoma and glioma. To study the effect of HS-SNVs on glioblastoma and glioma progression, we utilized the TCGA-GBM and LGG datasets and a GBM dataset from CGGA. Notably, higher expression of any of the HS-SNVs genes correlated with significantly better overall survival in the GBM-LGG and GBM datasets (Supplementary Figure S6A, S6B, S6C, and S6D). This highlights the prognostic significance of HS-SNVs in glioblastoma hypermetabolism.

### Known driver mutations are more frequent in hypermetabolic lesions

Driver genes play a critical role in glioblastoma evolution, and frequent mutations in *TP53, EGFR, PTEN,* and *TERT* genes have been well-defined (8,29,30). We analyzed single-nucleotide variants to observe the frequency and alterations of recognized glioblastoma drivers in hypermetabolic lesions. Intriguingly, TERT, TP53, FLG, SYNE1, and TTN were the most frequently mutated genes in zesto lesions at 90%, 60%, 30%, 40%, and 40%, respectively (Figure 5B). In contrast, only one–two out of 12 lesions were found to have alterations in kryo and akri lesions for the majority of driver genes. Additionally, the frequency of TERT and TP53 alterations in our cohort was in accordance with earlier studies that observed mutations in approximately 30– 50% of GBM samples. However, the frequency of EGFR alterations was lower (31,32), whereas FLG and SYNE1 frequencies were higher than those in previous reports (Supplementary Figure S6E) (32). Surprisingly, we observed only *EGFR* mutations in one kryo and akri lesion. Collectively, these analyses suggest that hypermetabolic lesions have a greater occurrence of recognized driver mutations than hypometabolic and peripheral lesions do. Such mutations may significantly influence the transformation of hypometabolic lesions into hypermetabolic ones.

### Hypermetabolic lesions exhibit a high frequency of fusion, omikli, chromothripsis, and chromoplexy events

Gene fusions are known to favor tumorigenesis, play an oncogenic role, and provide an opportunity for therapeutic targeting of glioblastoma and other cancer (33–35). We used whole-genome sequencing (WGS) and transcriptome analyses to compare and validate fusion events in our cohort. Across all samples, 182 fusion transcripts were detected, consisting of 26 deletions, 45 duplications, 31 inversions, and five translocations. Zesto lesions had a significantly higher number of these events than kryo and akri lesions (Figure 6A, Supplementary Figure S9, S10 and Supplementary Table S6). Some notable alterations in the zesto lesion were *PDGFRA-FIP1L1*, *PTPRZ1-MET*, *MTMR2-CEP57*, and *PDGFRA-TMEM165.* Moreover, frequent fusion events with duplications of *CAVIN4-MSANTD3-TMEFF1* and *CAVIN4-TMEFF1* in multiple kryo (50%) and akri (66.67%) lesions were observed (Supplementary Table S6). The occurrence of these fusion events was cross-validated with the STAR-Fusion tool using the RNA-seq data.

**Figure 6.**
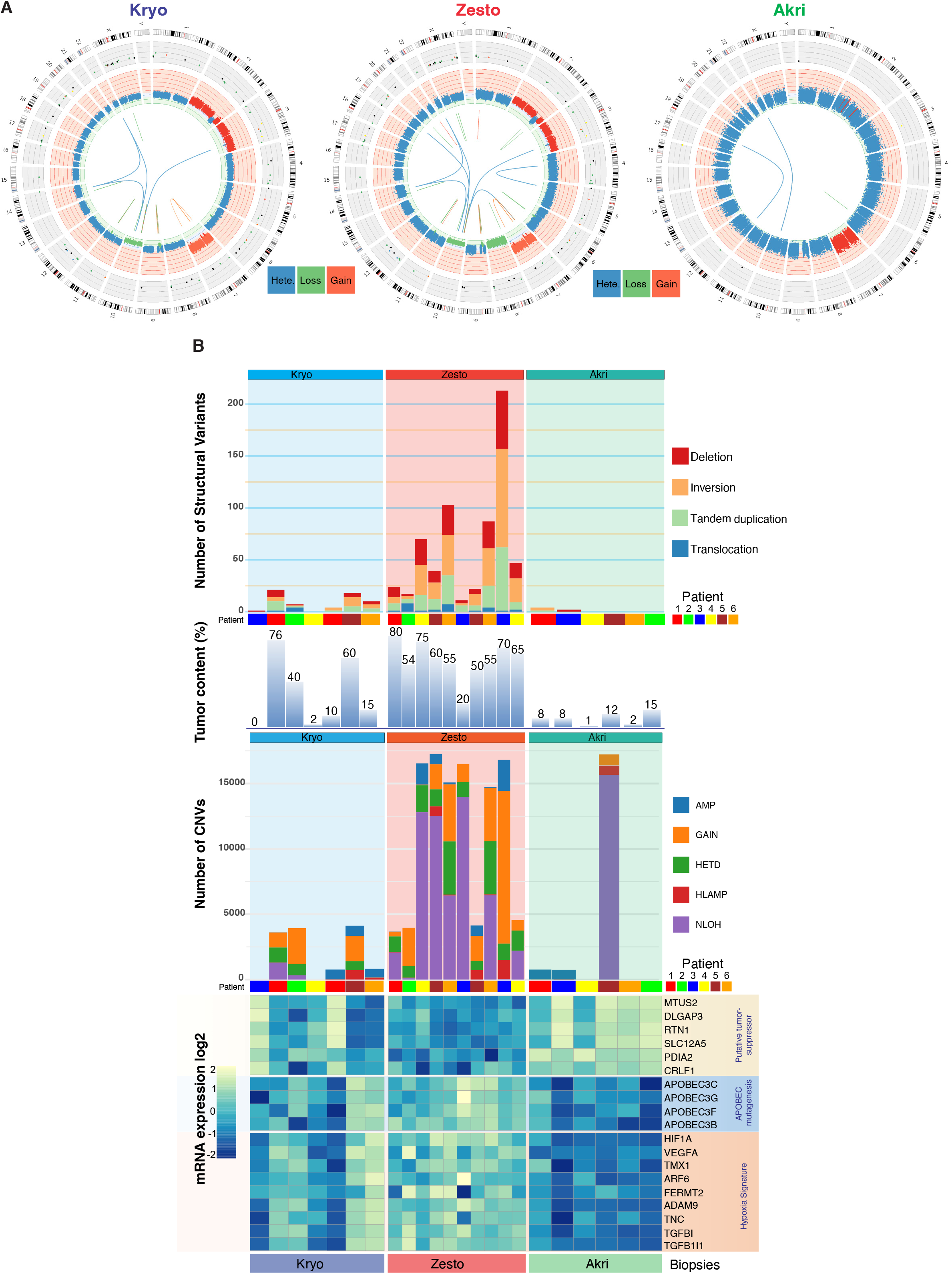
Occurrence of genomic rearrangements and chromothripsis events in the hypermetabolic lesions. **(A)** Representative scaled circos plots of the whole-genome of autologous lesions from a glioblastoma patient#5 showing genomic rearrangements. The outer ring shows the SNVs from the whole exome sequencing analysis, and the middle ring indicates the copy number variation from the whole exome sequencing, blue color represents heterozygosity, the red color indicates the gain, and the green indicates the loss of chromosomes. The innermost circle shows the structural variant analyzed by whole genome sequencing. **(B)** multi-omics global overview of structural variants in glioblastoma kryo (light blue background), zesto (light red background), and akri (light green background) lesions. The top bar plot shows the deletion, inversion, tandem duplication, and translocation in individual lesions. The second bar plot shows the tumor content in respective lesions, numbers represent the percentage of tumor content based on the whole exome sequencing estimation. The third bar plot indicates copy number variation such as (AMP) amplification, gain, (HETD) heterozygous deletion, (HLAMP) high-level amplification, (NLOH) neutral loss of heterozygosity, and (LOH) Loss of heterozygosity. The bottom heatmap shows the mRNA expression of putative tumor suppressor genes (*MTUS2, DLGAP3, RTN1, SLC12A5, PDIA2,* and *CRLF1*), APOBEC mutagenesis-associated genes, and hypoxia gene signature in the representative kryo, zesto, and akri lesions.

Diffuse hypermutations (omikli) (36), localized hypermutations (kataegis) (37), clustered chromosomal rearrangements (chromothripsis) (38), and complex DNA rearrangements (chromoplexy) (39) have been reported in glioblastoma and other cancer types. To assess these genomic phenomena, we analyzed WES and WGS data for all samples from our cohort. When comparing different lesions globally, we observed a higher frequency of translocations, duplications, tandem inversions, and deletion events in zesto lesions than in kryo and akri lesions (Figure 6B, top panel). Importantly, genomic events such as amplification, gain, heterozygous deletion, high-level amplification (HLAMP), and copy-neutral loss of heterozygosity (NLOH) events are also more frequent in zesto lesions. Interestingly, a high number of genomic alterations in zesto lesions anticorrelated with the expression of putative tumor suppressor genes (*MTUS2, DLGPA3, RTN1, CRLF1, SLC12A5,* and *PDIA2*) and positively correlated with *APOBEC* mutagenesis (*APOBEC3C, APOBEC3G*, *APOBEC3B* and *APOBEC3F*) along with hypoxia-related gene signatures (Figure 6B, bottom panel).

Moreover, we observed chromosomal shattering (chromothripsis) events in zesto lesions (4 out of 10 zesto) on chromosome 3 from patient-3 and chromosome 7 from patient-6 (Supplementary Figure S9). Indeed, we observed chromoplexy events often characterized by the involvement of DNA segments from multiple chromosomes, which were present in both zesto lesions from patient-6 (20% of hypermetabolic lesions). Kataegis (omikli, defined as diffuse mutational fog), characterized by localized hypermutation with hallmarks of a long range of C→T substitutions, was not observed in any biopsy. However, global base pair substitutions were dominated by C→T, followed by T→C in most hypermetabolic lesions, and hypometabolic lesions indicated the most frequent occurrence of omikli events (Supplementary Figure S10 B). These findings reflect the role of HS-SNVs. Specifically, the rational balance of MTUS2 and APOBEC3 expression may have a protective role in normal cell division, as the mutational burden is primarily associated with low mRNA expression levels of MTUS2, DLGAP3, RTN1, and SLC12A5. Altogether, elevated occurrences of fusion, omikli, chromothripsis, and chromoplexy events might be significant contributors to hypermetabolism in glioblastoma, potentially playing a crucial role in the shift from hypometabolic to hypermetabolic lesions.

## Discussion

Although many studies have reported complex tumor heterogeneity in glioblastoma, profound intratumoral, spatial, and genomic heterogeneity and its link to tumor metabolism used for glioma imaging in the clinical setting have been loosely elaborated (9,31,40–42). We demonstrated that hypermetabolic lesions identified by PET imaging have a higher frequency of genomic alterations than hypometabolic lesions, and that these alterations are associated with increased aggressiveness. These findings are supported by the following novel discoveries: (1) hypermetabolic lesions genetically evolve from hypometabolic lesions; (2) hypermetabolic lesions are spatially and genomically more diverse than hypometabolic lesions; (3) hypermetabolic lesion-specific CNVs and SNVs specifically alter (GPCRs, chloride transport, mitochondrial translational elongation and termination, and circadian rhythm) biological networks; and (4) downregulation of genes *MTUS2, DLGAP3, RTN1, SLC12A5, PDIA2, PHC3,* and *CRLF1* is associated with a higher frequency of fusion, omikli, chromothripsis, and chromoplexy events. Indeed, downregulation of this gene signature results in poor patient outcomes, indicating its potential prognostic value in glioblastoma. A schematic illustration of the genomic differences using multi-omics analysis is shown in Figure 7.

**Figure 7.**
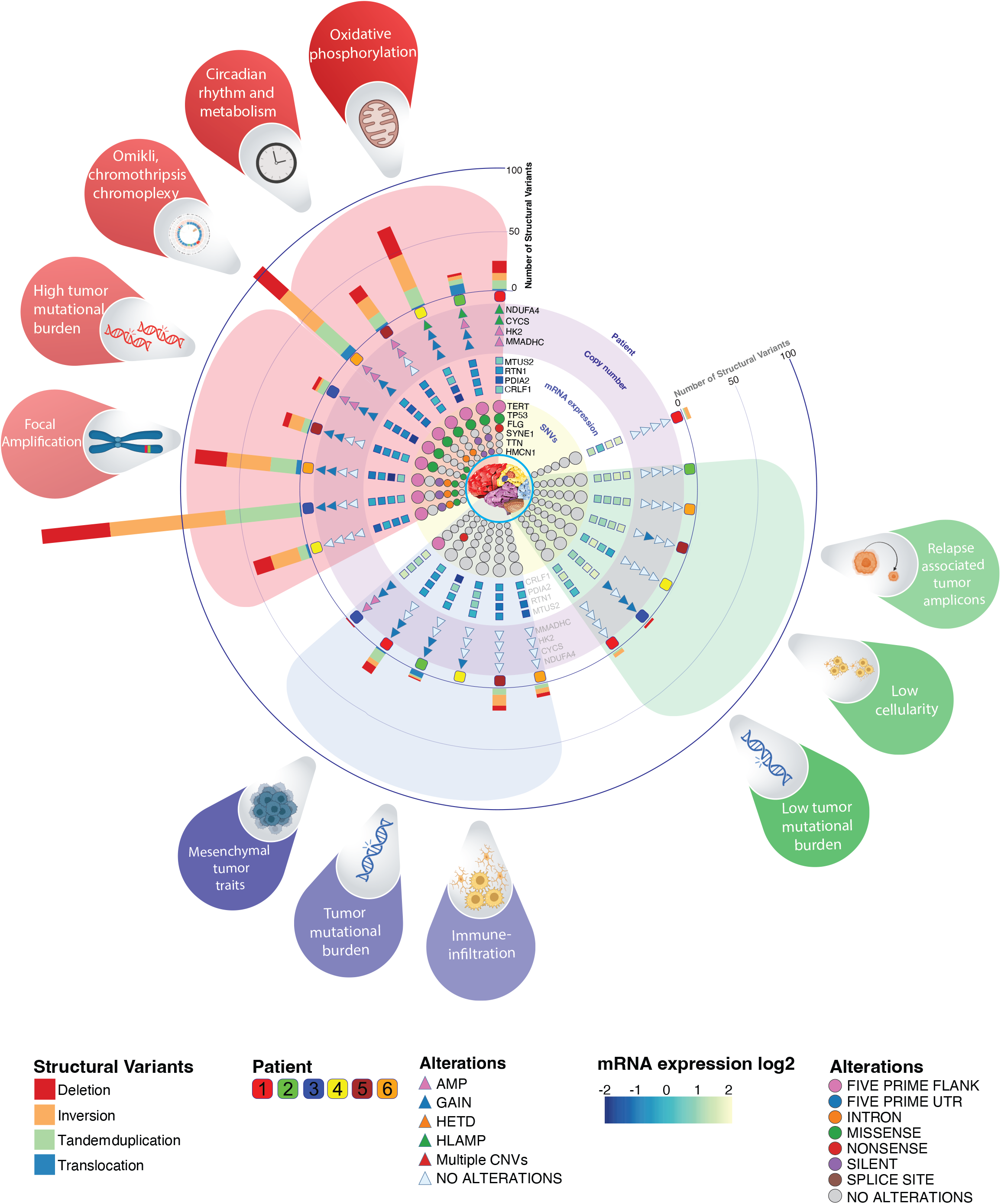
Graphical overview and comparison of the kryo, zesto, and akri lesions of glioblastoma patients: The innermost circle (in the round) shows single nucleotide variant analysis from whole exome sequencing data representing the top six most frequently mutated genes (*TERT, TP53, FLG, SYN1, TTN,* and *HMCN1*). The second circle (in the square) indicates the mRNA expression heatmap of putative tumor suppressor-like genes (*MTUS2, RTN1, PDIA2,* and *CRLF1*) in different lesion groups. The third circle (in the triangle) indicates the copy number alterations from whole exome sequencing analysis of representative genes (*NDUFA4, CYCS, HK2,* and *MMADHC*) from oxidative phosphorylation, gluconeogenesis, and amino acid metabolism pathways. The fourth circle (rounded rectangle) indicates the patient number. The stacked bar plot shows the number of structural variants based on whole-genome sequencing analysis. Abbreviations: (AMP) amplification, gain, (HETD) heterozygous deletion, (HLAMP) high-level amplifications, (NLOH) Neutral Loss of heterozygosity. Outer petals in blue shades (hypometabolic lesions: kryo), red shades (hypermetabolic lesions: zesto), and green shades (no metabolic activity lesions: akri) indicate the hallmarks of lesions. Inner petals in light blue, light red, and light green represent the kryo, zesto, and akri lesions sections.

Additionally, this study identified the intratumoral metabolic and genomic heterogeneity that regulates patient outcomes caused by HS-SNVs. However, we did not observe a very high mutational frequency of HS-SNV genes (*MTUS2, DLGAP3, RTN1, SLC12A5, PDIA2, PHC3,* and *CRLF1*) in other glioblastoma patients; however, it is tempting to speculate that the downregulation of HS-SNVs genes can be regulated by non-mutational epigenetic reprogramming (43,44) and may act as a switch for genomic abnormalities that lead to the evolution of hypermetabolic tumor lesions in glioblastoma.

Tumor evolution involves an intricate interplay between genetic and epigenetic factors as well as spatially diverse physiological microenvironments. Pre-mutational factors, specifically hypoxia and altered intracellular pH, are crucial for creating a conducive microenvironment for transformational changes, leading to the accumulation of genetic alterations. Importantly, a correlation between higher levels of hypoxia, pH, and a higher number of driver mutations has been observed in various types of cancer (45–47). We showed that the high expression of hypoxia gene signatures, such as *HIF1A*, *VEGFA*, *TMX1*, and *TGFBI*, among others, in hypermetabolic lesions positively correlates with APOBEC signature genes. APOBEC proteins, which cause the deamination of cytosine residues in both DNA and RNA, resulting in somatic mutations, RNA modifications, DNA breaks, or DNA demethylation (48,49), show a positive correlation with the hypoxia gene signature and tumor mutational burden in hypermetabolic lesions. In addition, cyclic hypoxia is known to cause a replication catastrophe, resulting in elevated APOBEC3B activity in bladder, breast, lung, and colorectal cancers (50). This result is consistent with our findings regarding hypermetabolic glioblastoma lesions. In particular, this study established a link between APOBEC expression, high tumor mutational burden, and the hypermetabolism of glucose and methionine in glioblastoma.

Previous studies have shown that *TERT* promoter and *TP53* mutations are among the earliest mutational events in glioblastomas and other cancer (51–53). Our analysis suggested that glioblastoma lesions with high metabolic activity commonly exhibit key driver mutations, particularly in genes such as TERT, NF1, TP53, FLG, SYNE1, and HMCN1, whereas hypometabolic lesions do not. Indeed, another study also suggested that TERT promoter mutations can be subclonal and may arise during later stages of tumorigenesis (30). Our findings indicate that there is a strong likelihood that driver mutations play a role in initiating transformation into hypermetabolic lesions. However, further research is needed to comprehensively understand the mechanisms through which driver mutations shape the increased metabolic capabilities of tumors. In daily clinical practice, the significance of this finding is important. For instance, the detection of a TERT promoter mutation in an IDH-wildtype astrocytic glioma is sufficient to establish glioblastoma diagnosis (54,55). For patients for whom needle biopsies are to be obtained for diagnostic purposes, it is advisable to extract such biopsies from hypermetabolic lesions guided by PET imaging.

Prior research focused on multifocal, multisectoral glioblastoma, as well as other cancer types, has shown a high level of mutational burden and diversity when compared to tumors located in close proximity (42,56,57). Additionally, a recent study revealed substantial intratumoral heterogeneity within gliomas while also drawing attention to unexpected homogeneity among brain metastasis samples (9). While some of our findings align with this research, earlier studies have not addressed the link between genomic alterations and tumor metabolism.

Glioblastomas are characterized by significant alterations in cellular metabolism, which serves as a prominent hallmark of these tumors (58,59). In glioblastomas, the Warburg effect plays a significant role in tumor metabolism. Glioblastoma cells exhibit a high rate of glucose uptake and preferentially utilize glycolysis as their primary energy source despite the availability of oxygen. However, in the absence of oxygen, the HIF-dependent hypoxic response regulates various cellular processes including metabolism, migration, angiogenesis, and differentiation (60). Our findings demonstrated that hypermetabolic glioblastoma lesions display a high number of copy number variations (CNVs) in genes involved in oxidative phosphorylation, gluconeogenesis, and amino acid metabolism pathways to meet the energy requirements of tumor cells. Additionally, we observed elevated expression of a hypoxic gene signature in hypermetabolic lesions, possibly attributable to intense cell cycling that leads to oxygen and nutrient deprivation. In addition, our results affirm the findings of a study by Wu et al. (6), which suggested that hypoxia is involved in inducing the natural evolutionary signature of the tumor by HIF1A.

Altogether, our multi-omics analysis established that human glioblastoma with different metabolic lesions demonstrates substantial genomic heterogeneity and continuous tumor evolution. Hypermetabolic lesions in glioblastomas evolve and exhibit a more advanced mutational profile than hypometabolic lesions, posing a challenge for effective treatment. In light of our findings, we recommend that surgeons obtain biopsies from hypermetabolic lesions in glioblastomas identified through PET and MRI imaging for diagnostic purposes. To attain a more profound understanding of the tumor’s mutational traits and its complex biology, future endeavors should involve targeted sampling coupled with multiomics profiling. This is critical for improving personalized therapeutic strategies for glioblastoma treatment.

## Methods

### Glioblastoma patients and clinical annotation

All the patients included in this study were histologically diagnosed with glioblastoma. With a focus on performing and integrating multi-omics analysis, glioblastoma patients were only selected if the tumor tissue, normal tissue (D400 mg), and blood specimens (D3 ml) were sufficient to perform the multi-omics analysis. In total, 23 fresh frozen glioblastoma patient tissues and 6 blood samples were collected at the Department of Neurosurgery at Odense University Hospital (Odense, Denmark) after informed ethical consent from the patients was obtained and approved by the Regions Scientific Ethical Committee of Southern Denmark (S-20140214). Clinical data, such as age, sex, tumor location, tumor size, diagnosis, IDH, and MGMT methylation status, were obtained from the Department of Pathology. Fresh tissue specimens from patients were collected during surgical resection and snap-frozen in liquid nitrogen for 45 min before being stored at −80°C. The AllPrep DNA/RNA Mini Kit (Qiagen #80284) to extract DNA and RNA from each sample. The DNA was measured using a Qubit fluorometer. RNA quality and quantity were measured using an Agilent 2100 Bioanalyzer.

### MRI and PET imaging

Patients included in this study were histologically diagnosed with glioblastoma based on a positive needle biopsy. The patients were >48 years old and scheduled for surgery the day after FDG-PET/MET-PET/MRI. Conventional MRI was performed as a standard procedure, and positron emission tomography (PET) with 2-deoxy-2-[fluorine-18] fluoro-D-glucose (^18^F-FDG), an analog of glucose, and L-[methyl-^11^C] methionine (^11^C-MET), an analog of methionine, was performed separately on PET/CT scanners at the Department of Radiology, Odense University Hospital, Odense, Denmark. MRI was performed using the Medtronic Stealth Neuronavigation System. T1w and T2w structural MRI images were acquired at an isotropic resolution of 0.7 mm. The T1w parameters were repetition time (TR) =2,400 ms, echo time (TE) =2.14 ms, inversion time (TI) =1,000 ms, flip angle (FA) =8°, while the T2w parameters were: TR=3,200 ms, TE=565 ms. For rs-fMRI imaging, each of the two rs-fMRI sessions consisted of two scans acquired in opposite phase encoding directions (left-to-right and right-to-left) with a voxel size of 2 mm^3^ and parameters: TR=720 ms, TE=33.1 ms (61).

### Stereotactic biopsy isolation

To plan lesions from preoperatively defined areas in the tumor and its vicinity, MRI was co-registered with 11C-MET-PET and 18F-FDG PET. Areas of interest were predefined, and during surgery, craniotomy was performed to ensure that the dura was not accidentally lesioned. A neuronavigational inaccuracy <1 mm was considered acceptable. To avoid brain shift, 3-dimensional stereotactic image-guided needle lesions were harvested prior to dural opening. Regions with low uptake of 18F-FDG and 11C-MET were classified, and the lesions were labeled as metabolic kryo, high uptake as Zestos, and no uptake as akri in six glioblastoma patients. In one patient (patient #1), we observed and isolated a biopsy specimen surrounding the main tumor mass, called a tumor satellite. The average distance between all lesions was measured using a navigation system. The GE AW Server platform (Dexus) software was used to analyze, visualize, and co-register the FDG-PET/MET-PET/MRI scans. The normal surgical time for each patient was 90–120 min, and the stereotactic biopsy procedure was performed for an additional 60 min.

### Hematoxylin and Eosin staining

A representative small part of the biopsies was formaldehyde-fixed, embedded in paraffin, and cut into 3µm sections. To visualize cellularity, the slides were stained using the standard protocol for Hematoxylin and Eosin staining. The stained sections were scanned qualitatively using a NanoZoomer 2.0-HT slide scanner and the NDP viewer software (Hamamatsu).

### Whole Exome sequencing data analysis

1000ng of DNA was used to prepare the library using the KAPA Target Enrichment Kit (Roche) for exome sequencing. Raw reads were aligned and duplicate-marked using the DRAGEN Bio-IT Platform (Illumina). Somatic variant calling was performed using VarScan v. 2.3.4. Only bases with a quality score of at least Q20 were considered. Variant annotation and filtration were performed using VarSeq (Golden Helix) software. We used the following criteria to identify somatic mutations derived from the exome data: genotype quality (QC) >100, read depth of >100x and VAF >5% in one of the tumor samples and VAF < 2%, and allele count of < 3 variant counts in the normal sample. CNV calling, filtering, and visualization were performed using VarSeq (Golden Helix). The following criteria were used for copy number variants: number of exons > 5 and average coverage >100x. All sequencing procedures were performed in a single run. One NovaSeq S4 lane was used for RNA-Seq, two lanes were used for exomes, and one lane was used for genome sequencing (2*150bp).

### Whole Genome sequencing and quantification of structural variant data analysis

We used 200ng of DNA to prepare the library. To achieve 25X coverage for all samples, whole-genome libraries were subjected to 150 bp paired-end sequencing on the Illumina NovaSeq 6000. The FASTQ files resulting from whole-genome sequencing were pre-processed according to GATK’s best practices to obtain analysis-ready BAM files. BRASS (The Breakpoint Analysis tool https://github.com/cancerit/BRASS) was used to identify structural variants and TitanCNA (10) was used to identify copy number variants (CNVs).

### RNA Sequencing

mRNA libraries were generated from 10 ng of RNA from each sample using TruSeq® RNA Library Prep for Enrichment (20020189) and TruSeq® RNA Enrichment (20020490) library prep kits, following the manufacturer’s protocol. The libraries were sequenced on an Illumina NovaSeq 6000 System high-output flow cell (2 × 150bp), with one lane of an S4 flow cell used for transcriptome sequencing of all 23 biopsies. A total of 2,568,575,326 reads were obtained and aligned to the human reference genome (GRCh38) using STAR. Duplicate reads were removed, and the remaining reads were sorted using SamTools. Gene expression counts were generated for each sample using featureCounts in the R subread package. The resulting counts were normalized to counts per million (CPM) and subjected to trimmed mean M (TMM) normalization using edgeR. During the TMM normalization step, genes with counts below a certain CPM cut-off were filtered out to reduce noise. On an average, approximately 75% of the reads were uniquely mapped.

### Tumor content analysis

Tumor content was estimated using a set of CNV tools: TitanCNA (10) (on WGS and WES data), AscatNGS (62) (on WGS and WES data), Sequenza (63) (on WES data), and Battenberg (64) (on WGS data) using the default settings. Together with the computed estimates by these tools, we also analyzed plots of one sample’s variant allele frequencies (VAFs) from the WES data analysis against the VAFs from another sample to identify the center of heterozygous variants, which, in an ideal scenario, corresponds to half of the tumor content present in each sample. A combination of these two approaches was used to estimate the best possible tumor content for each sample.

### Public datasets and analysis tools

Normalized gene expression, patient survival, gene correlation, and mutational analyses in low-grade glioma (LGG) and glioblastoma multiforme (GBM) were performed on projects from TCGA, CGGA, the Ivy Glioblastoma Atlas Project (Ivy GAP) dataset, and GEPIA (65); data were obtained from the GlioVis website (http://gliovis.bioinfo.cnio.es/) (16). Oncoscore analysis of the HS-SNVs gene was performed using the Oncoscore web tool (24) (https://www.galseq.com/next-generation-sequencing/oncoscore-software/). OncoGrid analysis (Figure 5D) was performed using the Genomic Data Commons Data Portal (National Cancer Institute) and the TCGA-GBM dataset (https://portal.gdc.cancer.gov/). Aneuploidy score analysis was performed using the cBioportal tool (https://www.cbioportal.org/) by checking the availability of all glioblastoma datasets with genomic analysis (27,28) using default settings. Single-cell sequencing data analysis was performed using Loupe cell browser software from 10X Genomics. Raw data (Male, Glioblastoma patient) were downloaded from the 10X Genomics website (https://www.10xgenomics.com/) with consent and permission. Single gene ontology enrichment analysis of NMU was performed using the Gliovis tool (http://gliovis.bioinfo.cnio.es/) and the TCGA-GBM datasets. Briefly, we used the path for differential expression tab/gene ontology tab/dot plots using default settings.

### Analysis of protein-protein interactions between zesto biopsy-specific CNVs

To study the physical interactions between potential driver genes, we used the top 470 gene lists from zesto biopsy CNVs as input gene lists for the STRING database (66). As shown in Figure 4E, we used all zesto-biopsy-specific SNVs with 21 genes as an input list. A minimum required interaction score of 0.700 was used to achieve high-confidence interaction predictions.

### In silico projection of SNVs impact

We used dbNSFP (22) v4, a database for the functional prediction of potential non-synonymous single-nucleotide variants (nsSNVs) in the human genome. dbNSFP uses functional prediction voting (six votes) to predict the impact of the SNV. SNVs identified from zesto-specific lesions were used as inputs for dbNSFP v4 tools. Deleterious SNVs were predictions that were possibly or probably damaging (Polyphen2) or neutral, medium, or high (Mutation Assessor).

### Mutation burden analysis

SNV and indel burden analyses were performed using the formula for the number of SNVs identified in the WES data by the Varscan tool and divided by the total area of the exome analyzed.

## Data availability

The expression matrix for mRNA sequencing is provided in the Supplementary Data. This study was performed with permission from the Danish law, and informed consent was needed for such research. However, the publication only contains aggregated results, and no personal data are disclosed in this manuscript. The gene expression data can be obtained from the NCBI GEO database with accession number GEO: XXXXXXX. (Accession number awaited). The whole-genome and exome sequencing data used in this work are person identity-sensitive, and to abide by the rules, we are prohibited from making them publicly available according to Danish legislation. However, access to raw genomic data for the research community can be provided upon request with an agreement to protect the confidentiality of individuals. This request can be written as.

### Statistical analysis and data visualization

Data analysis and visualization were performed using GraphPad Prism (https://www.graphpad.com/scientific-software/prism/), R software (http://www.r-project.org), Biorender (https://biorender.com), and Adobe Illustrator-2023 (https://www.adobe.com/products/illustrator.html). Unless otherwise specified, all statistical analysis tests were two-tailed, and a P or FDR value of <0.05 was considered statistically significant. Where applicable two-sample t-tests, and linear models were used, and the resulting P values are reported in the figure legend.

## Acknowledgments

We thank all patients and their relatives who agreed to participate in this study at Odense University Hospital, Odense, Denmark. The helpful assistance of operating room nurses at the Department of Neurosurgery, Odense University Hospital, Denmark, is gratefully appreciated. We thank Helle Wohlleben, Nicolai Stilling Schou, Eva Rahtkens Norup, and Nadine Margaretha Hammouda for their assistance with the laboratory. We thank Clara Rosa Levina Oudenaarden, Henning Boldt, Justin D. Lathia, Guido Reifenberger, Signe Regner Michaelsen, Dylan Scott Lykke Harwood, and Kartikey Saxena for their useful discussions and valuable comments. Part of this project was supported by the Regions of Southern Denmark Research Funds and the Novo Nordisk Foundation (NNF19OC0058427).

## Author’s Disclosures

All authors declare no competing interests.

## Authors’ Contributions

**A. Anand:** Conceptualization, resources, data analysis and interpretation, investigation, visualization, funding acquisition, writing of the original draft, writing– review, and editing.

**J. K. Petersen:** Conceptualization, methodology, pathology information.

**L. v. B. Andersen:** formal analysis WES-WGS, investigation, visualization.

**Mark Burton:** formal analysis of transcriptome, investigation, and visualization.

**Martin Jakob Larsen:** formal analysis WES-WGS, investigation, visualization.

**Christian Bonde Pedersen:** Stereotactic biopsy and surgical removal of biological materials.

**Frantz Rom Poulsen:** Conceptualization, stereotactic biopsy, and surgical removal of biological materials.

**Peter Grupe:** Acquisition of brain imaging data and analysis.

**Torben A. Kruse:** Conceptualization, supervision, funding acquisition, project administration, writing, review, and editing.

**Mads Thomassen:** Conceptualization, supervision, funding acquisition, project administration, writing, review, and editing.

**Bjarne Winther Kristensen:** Conceptualization, pathology analysis, supervision, funding acquisition, methodology, project administration, writing, review, and editing. All the authors have read and approved the final manuscript.

## Notes

**Funding:** This study was funded by the Danish Cancer Society, the Region of Southern Denmark Research Funds, and the Novo Nordisk Foundation.

### Competing Interest Statement

The authors have declared no competing interest.

