## Supplemental figures S1 to S12 for "Insight into spatial intratumoral genomic evolution in glioblastoma"

**Supplementary figure legends:**

**Supplementary Figure S1:**
**(A)** Tumor mutational allele frequency analysis by whole-exome sequencing in autologous kryo (hypometabolic tumor lesion) on the x-axis and zesto (hypermetabolic tumor lesion) on the y-axis from two glioblastoma patients with low tumor content in kryo. Zesto and kryo-specific variants are marked respectively with red and blue. Data points represent mutations the red dotted line represents perfect uniformity.
**(B)** Representative 3-dimensional geometric localization of lesions in the coronal, axial, and 3D view from a single patient. The yellow dashed line represents the tumor area, and the green circle is the point of needle insertion to pick up lesions.
**Table 1.** Shows the demographics of glioblastoma multiforme (GBM) patients involved in the study.

**Supplementary Figure S2:** MRI, [$^{18}$F] FDG-PET, [$^{11}$C] MET-PET, and co-registration images from all glioblastoma patients showing the location of the hypometabolic (blue circle), hypermetabolic (red circle), and akri (green circle) lesions. Blue intensity represents the lowest, and red represents the highest accumulation of radiotracers.

**Supplementary Figure S3: Spatial genomic heterogeneity in autologous lesion in glioblastoma** Merged transverse MRI, concordant FDG-PET, and MET-PET images exhibiting different geo locations of lesions and textboxes represent key lesion-specific chromosomal alterations in the patients.

**Supplementary Figure S4: Single-cell transcriptomic profiling highlights proliferating tumor cell clusters and co-expression of hypermetabolic lesion-specific genes involved in CNVs events.**
**(A)** Uniform Manifold Approximation and Projection (UMAP) plot of the location-averaged transcriptome for all cells (5604 cells) from a glioblastoma patient (male 57 years) obtained from 10x Genomics. Different cell types are colored by cluster (top left), the proliferative tumor cluster (based on MKI67 expression) is marked in the UMAP plot, and the top 10 highly expressed genes in the proliferative tumor cluster.
**(B)** The expression of genes (*DLGPA3, NMU, PAICS, CEP135, PPAT,* among other genes) involved in zesto lesion-specific CNV events shows a cellular colocalization with proliferative tumor cell gene signature.
**(C)** mRNA expression and copy number alteration of zesto lesion-specific cancer driver genes in the glioblastoma and low-grade glioma samples from the TCGA dataset (Homdel: Homozygous deletion, HetLoss: Loss of heterozygosity).
**(D)** Overall patient survival analysis of glioblastoma (163 patients) and low-grade glioma (518 patients) based on the high and low expression of zesto lesion-specific signature genes. Signature genes are mentioned in the lower left corner of plots, data analyzed by the GEPIA-2 web tool.

**Supplementary Figure S5: MTUS2 as a potential tumor suppressor, regulator of metabolism, and cell proliferation in glioblastoma:**

**(A)** Single-cell mRNA expression pattern of metabolic zesto lesion-specific SNV genes in a glioblastoma patient from 10X genomics dataset. The top left UMAP plots specify the major cellular subtype, and the rest of the UMAP plots indicate the expression or no expression of the zesto lesion-specific SNV genes. Most SNV genes overexpressed in the glioblastoma dataset are expressed in tumor and endothelial cell clusters. MTUS2, DLGAP3, RTN1, CRLF1, and PDIA2, have the lowest expression in the majority of cell types.

**(B)** mRNA expression of putative tumor suppressor genes MTUS2, DLGAP3, and RTN1 in glioblastoma and low-grade glioma dataset, statistical significance ($P<0.01$) were analyzed by one-way analysis of variance (ANOVA) and is indicated with an asterisk (analyzed by GEPIA2).

**(C)** MTUS2 mRNA expression negatively correlates with glucose metabolism-associated genes such as HK2, GALM, GAPDH, ENO1, HK3, PKM, and LDHA mRNA expression (TCGA-glioblastoma dataset: CBioportal).

**(D)** Protein abundance of MTUS2, SLC12A5, and RTN1 negatively correlates with cell proliferation/cycle-associated proteins KI67 and TK1. The data were analyzed using the cBioPortal web tool.

**(E)** zesto lesion-specific genes (MTUS2, DLGAP3, RTN1, CRLF1, and PDIA2) mRNA expression shows a positive correlation with each other, text box indicating the Spearman's and Pearson correlation and p-value in the bracket (TCGA-glioblastoma dataset: CBioportal).

**Supplementary Figure S6: Low expression of putative tumor suppressor signature genes associated with poor survival of glioma and glioblastoma patients.**

**(A)** Kaplan– Meier overall survival estimates of glioblastoma and low-grade glioma (667 patients) patients according to the high and low expression of genes.

**(B)** Kaplan Meier plot displaying CGGA dataset overall glioblastoma (WHO grade IV, 220 patients) survival distributions of patients stratified and colored by high and low expression of genes. Data was analyzed by the GlioVis web tool.

**(C) and (D)** gene expression and overall survival of Polyhomeotic Homolog 3 (*PHC3*) and Collagen Type XVIII Alpha 1 Chain (*COL18A1*) in TCGA (The Cancer Genome Atlas) IDH-wildtype glioblastoma (GBM), IDH-mutated 1p19q co-deleted oligodendroglioma (O) and IDH-mutated 1p19q non-co-deleted astrocytoma. *PHC3* and *COL18A1* were found to be mutated in hypermetabolic lesion.

**(E)** OncoGrid from WGS analysis performed on TCGA-GBM samples shows genetic alterations of the selected driver genes from the hypermetabolic lesion-specific gene signature.

**(F):** NMUR1 (receptor for *NMU*) mRNA expression indicates the expression primarily on the microvascular proliferation region, data analyzed by IvyGAP Dataset. NMU (ligand for NMUR1) is secreted by tumor cells, over-expressed in zesto biopsies.

**Supplementary Figure S7: The hypermetabolic (zesto) lesion exhibits micro-amplification and deletion of the chromosomal region that could potentially enhance tumor aggressiveness.**

**(A)** Global copy number variation in autologous lesions from glioblastoma patients (#2, #5) as identified using the whole exome sequencing workflows by the Varseq analysis tool. The x-axis depicts the genomic location, and the Y-axis indicates the log ratio of copy number. Arrows indicating lesion-specific copy number alterations.

**(B)** mRNA expression of genes located in the amplicon of patient-2, Chromosome-9p, marked with a red arrow.

**(C)** mRNA expression of genes located in the deleted region of the zesto lesion (Patient#2, Chromosome#9p, marked with red arrow), genes are arranged in the chromosomal order to show the pattern of CNV and gene expression difference between autologous zesto, kryo, and akri lesions.

**(D)** mRNA expression of *CDKN2A*, *CDKN2B*, and *MTAP* gene located in the zesto's deleted region (Patient#5, Chromosome-9p, marked with red arrow).

**Supplementary Figure S8:** Global copy number variation in autologous lesions from glioblastoma patients (3, 4, and 6) as identified using the whole exome sequencing workflows by the Varseq analysis tool. The x-axis represents the genomic location, and the Y-axis indicates the log ratio of copy number. Arrows indicate lesion-specific copy number alterations found in the lesions.

**Supplementary Figure S9: Hypermetabolic tumor lesions have higher genomic instabilities:** Circos plots of the genome of autologous lesions from different glioblastoma patients showing genomic rearrangements and chromothripsis (marked with a red arrow in zesto lesions from patients 3 and 6). The outer ring depicts the SNVs from the whole exome sequencing analysis. The middle ring indicates the copy number variation from whole exome sequencing, the red color indicates the gain, the green indicates the loss of chromosomes, and the blue shows heterozygosity. The innermost circle shows the structural variation analyzed by whole genome sequencing. The red arrow marks chromothripsis events, and chromoplexy events are marked by a dark grey arrow.

**Supplementary Figure S10: Hypermetabolic tumor lesions have higher genomic instabilities:**

**(A)** Circos plots of the genome from the zesto, kryo, and akri from patient 6. The outer ring depicts the SNVs from the whole genome sequencing analysis.

**(B)** Substitution plots in the respective lesions, indicating the highest substitution events were found in zesto compared to kryo and akri, C to T substitution was the most dominant and followed by T to C substitutions.

**(C)** Bar plots showing the quantification of deletion, insertions, and global rearrangements in different lesions from whole genome sequencing analysis.

**Supplementary Figures S11 and S12: Hypermetabolic tumor lesions exhibit a greater degree of genomic instability.**

A global overview of B- allele frequency analysis on all 29 samples to highlight the major occurrence of a CNV, such as a deletion, duplication, or loss of heterozygosity (LOH), in the whole genome region. **(Figure S11 for patients 1,2 and 3 and Figure S12 for patients 4, 5, and 6)**

Supplementary Figure S1

A

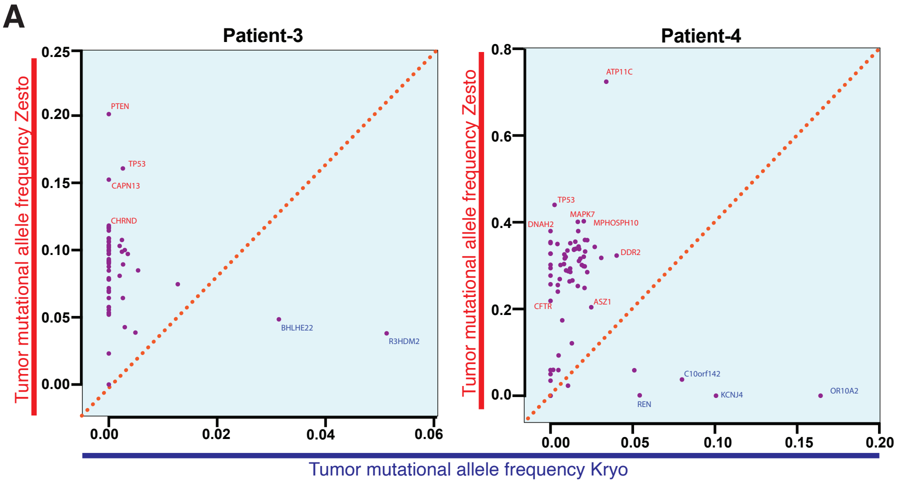


B

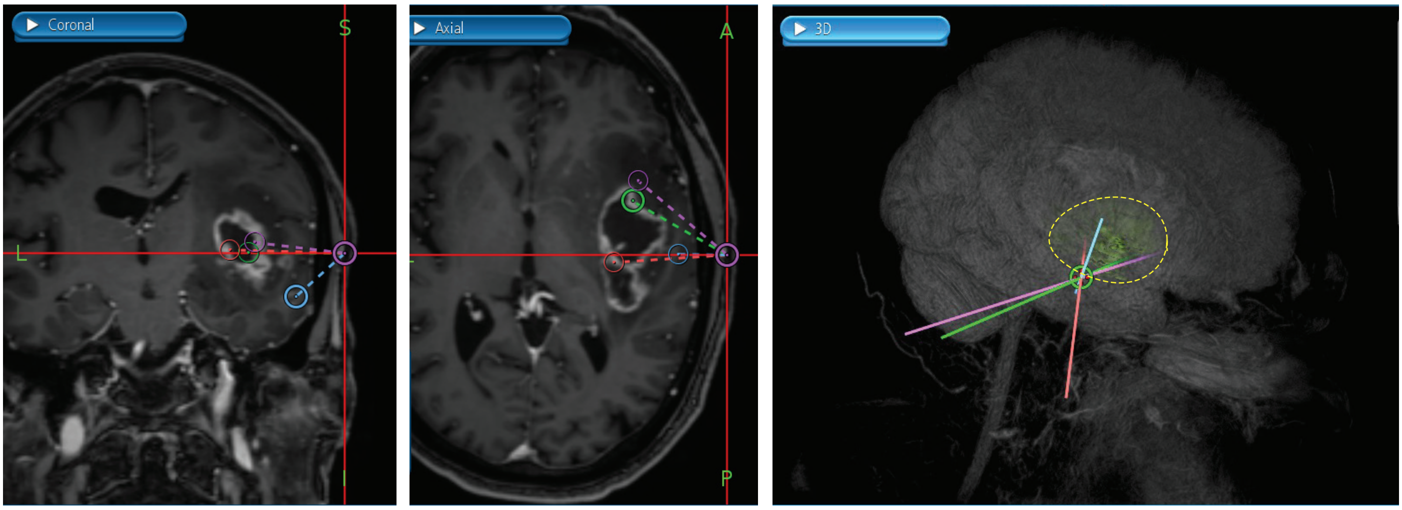


#### Table 1. Demographics of GBM patients

| Patient number | Age | Sex | Diagnosis | MGMT Promoter Methylation | IDH | WGS samples | WES samples | Transcriptome samples | Prior Treatments |
|---|---|---|---|---|---|---|---|---|---|
| 1 | 68 | male | GBM | unmethylated | wildtype | 4 | 5 | 4 | No |
| 2 | 74 | male | GBM | methylated | wildtype | 3 | 4 | 3 | No |
| 3 | 48 | male | GBM | methylated | wildtype | 4 | 5 | 4 | No |
| 4 | 76 | male | GBM | unmethylated | wildtype | 4 | 5 | 4 | No |
| 5 | 72 | female | GBM | unmethylated | wildtype | 4 | 5 | 4 | No |
| 6 | 72 | male | GBM | methylated | wildtype | 4 | 5 | 4 | No |

Supplementary Fig. S2

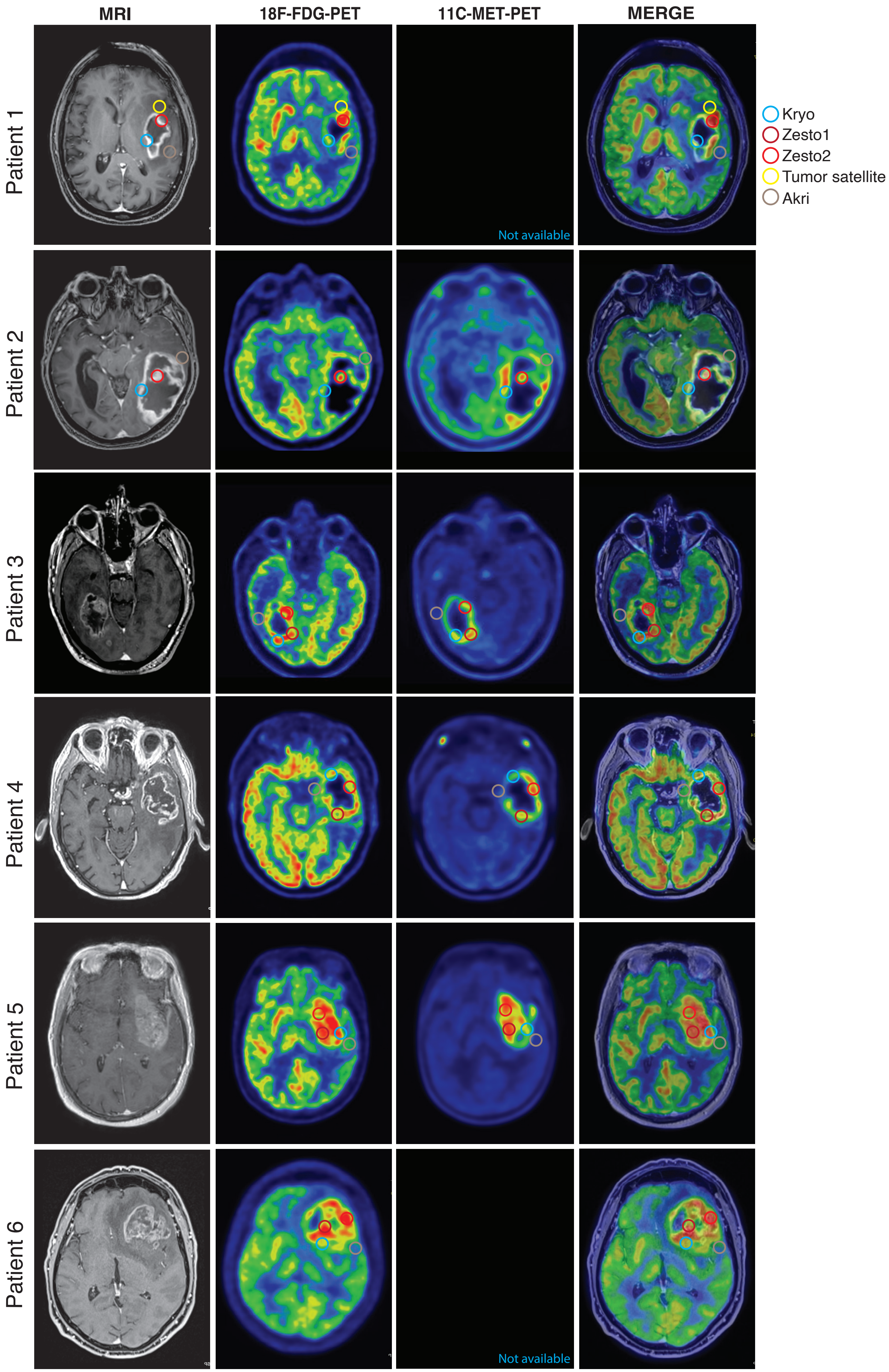

### Supplementary Figure 3

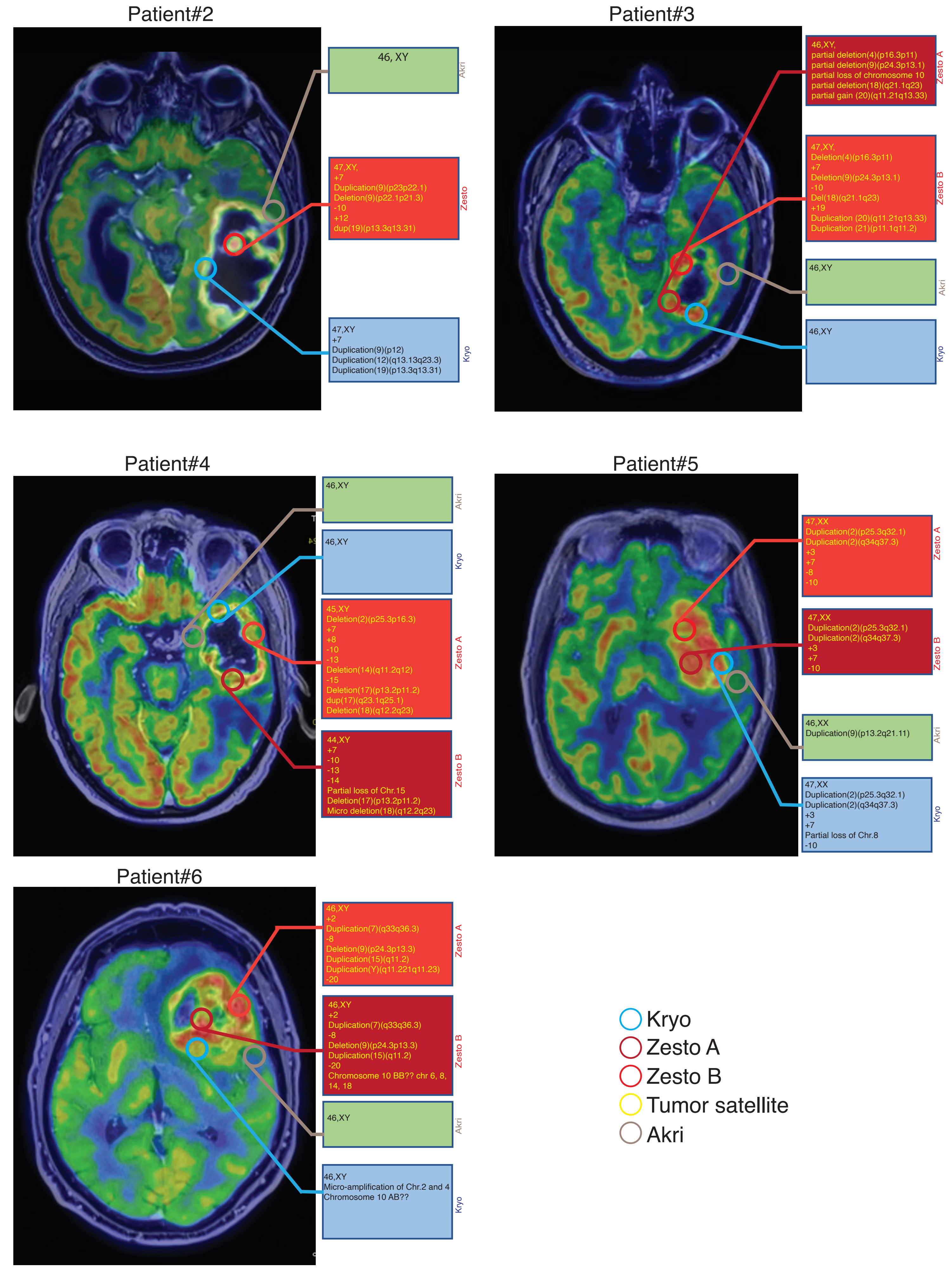

### Supplementary Figure 4

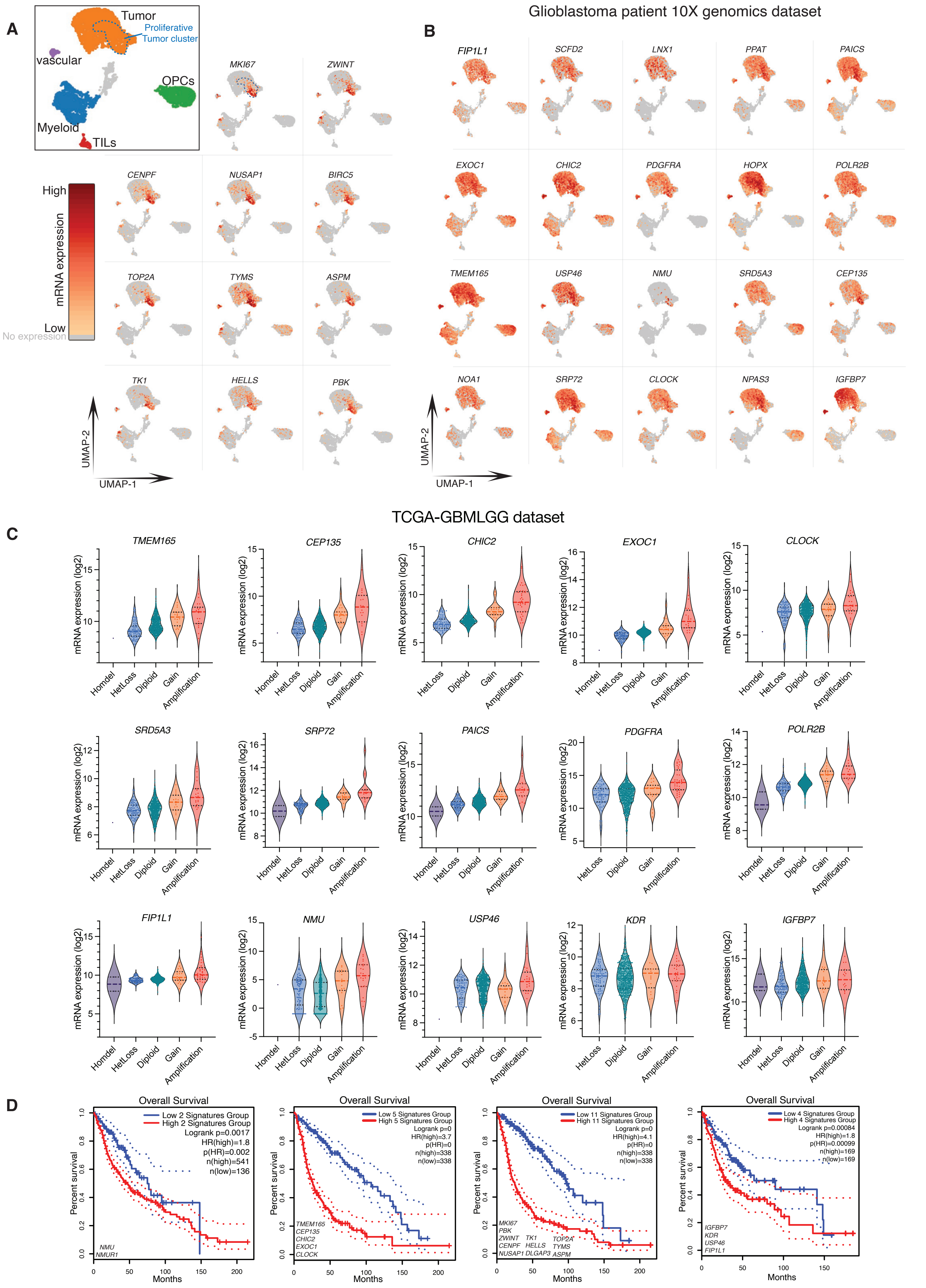

### Supplementary figure S5

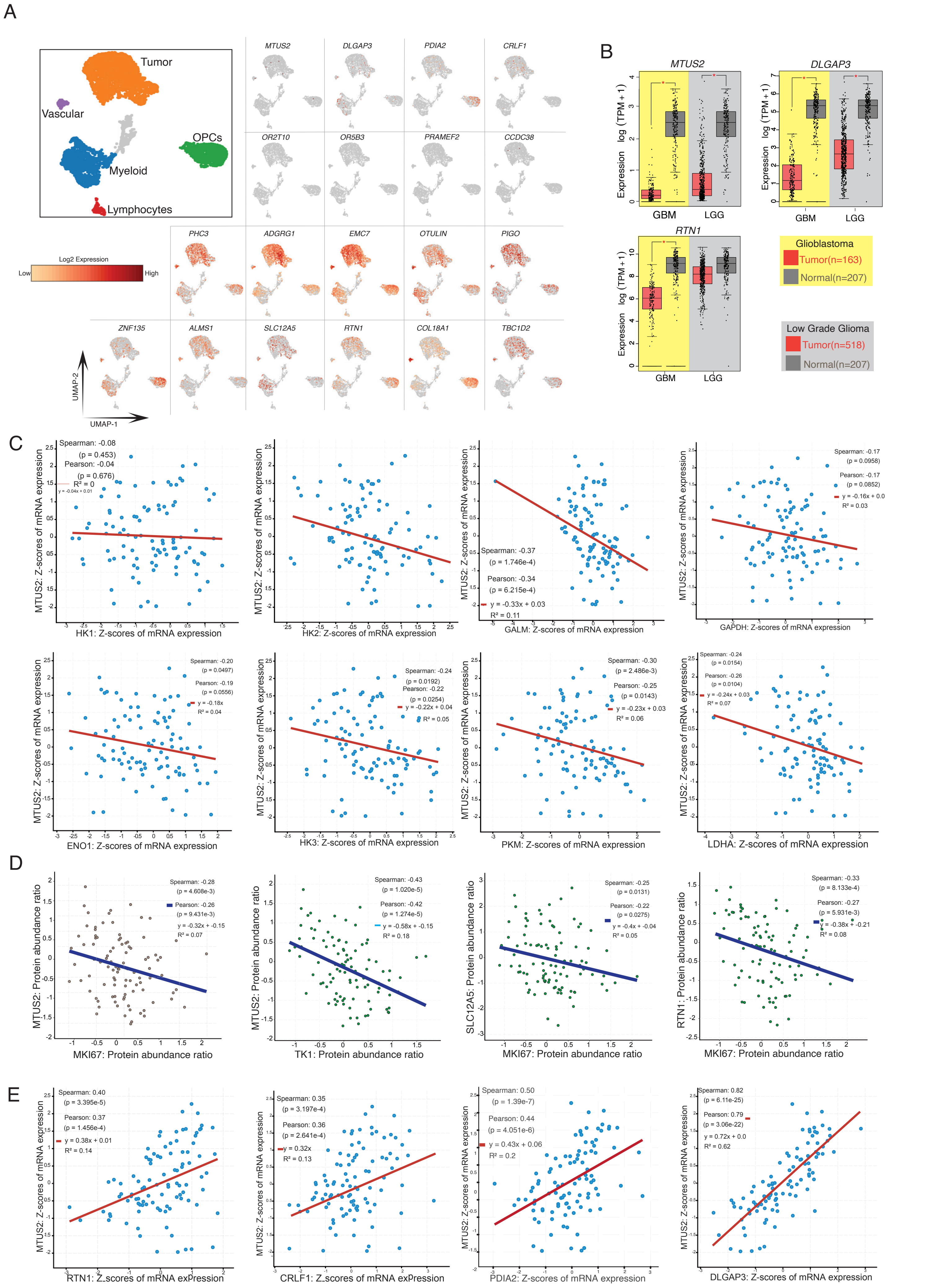

### Supplementary figure S6

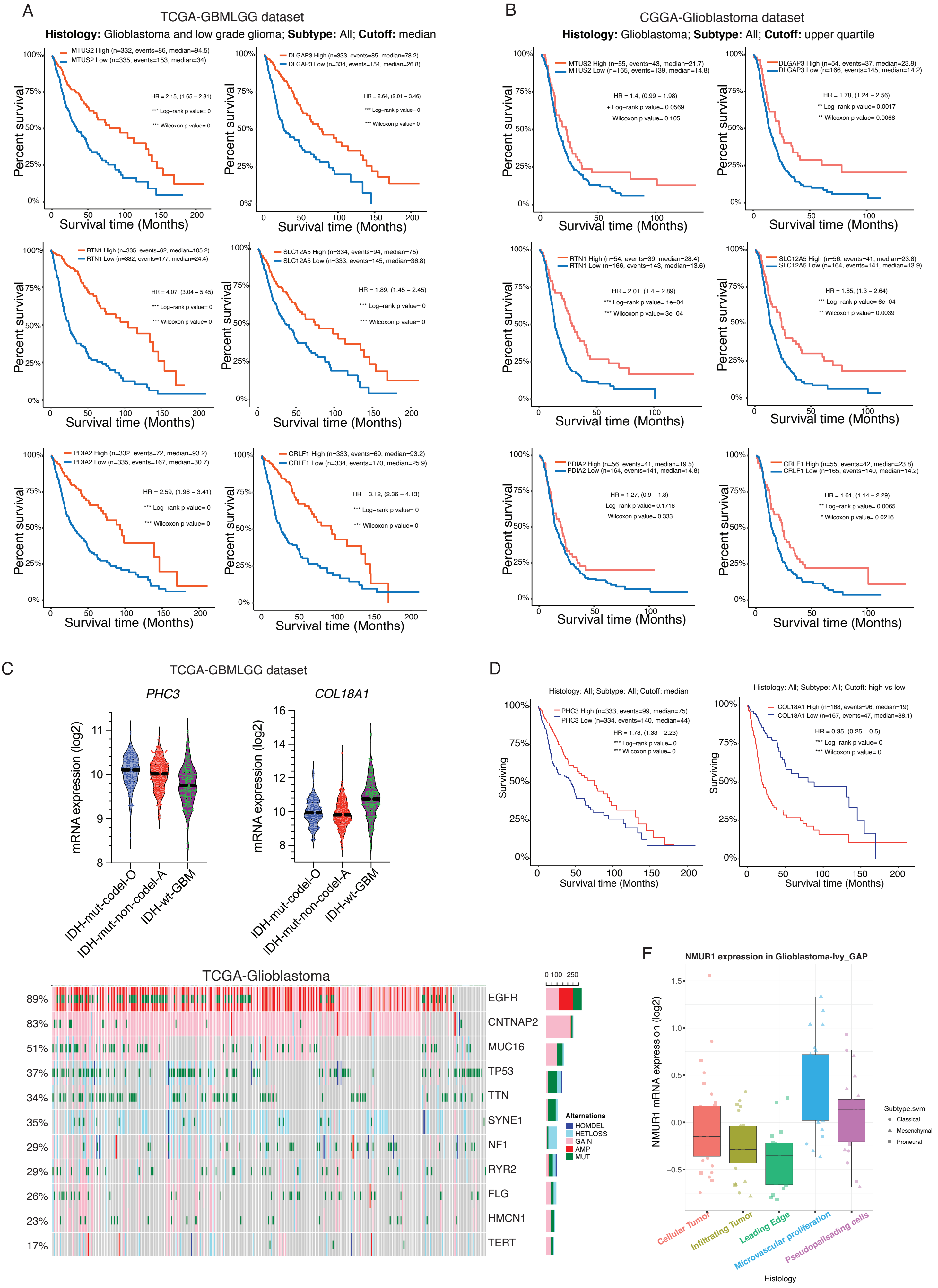

Supplementary Fig. S7

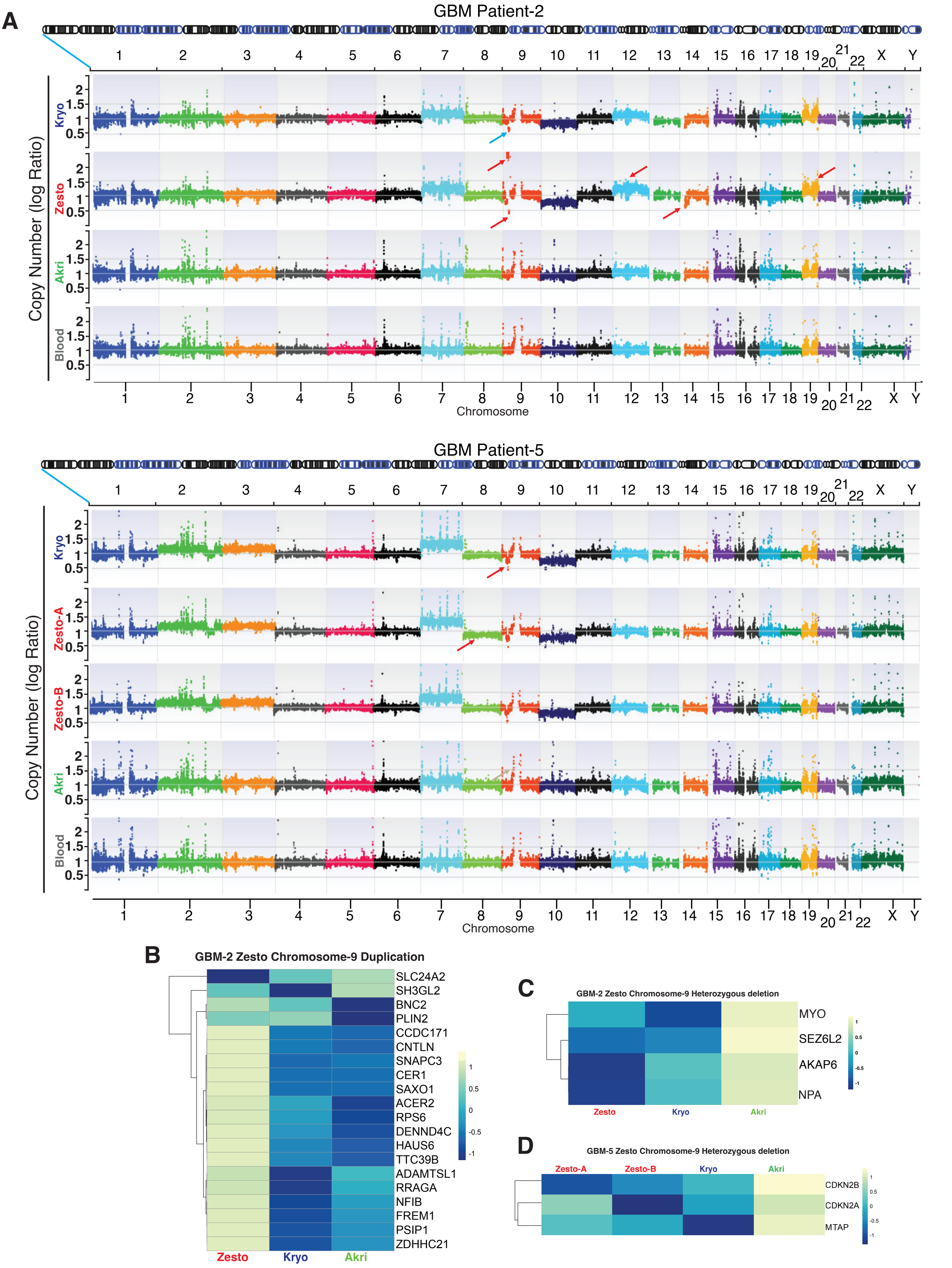

Supplementary Fig. S8

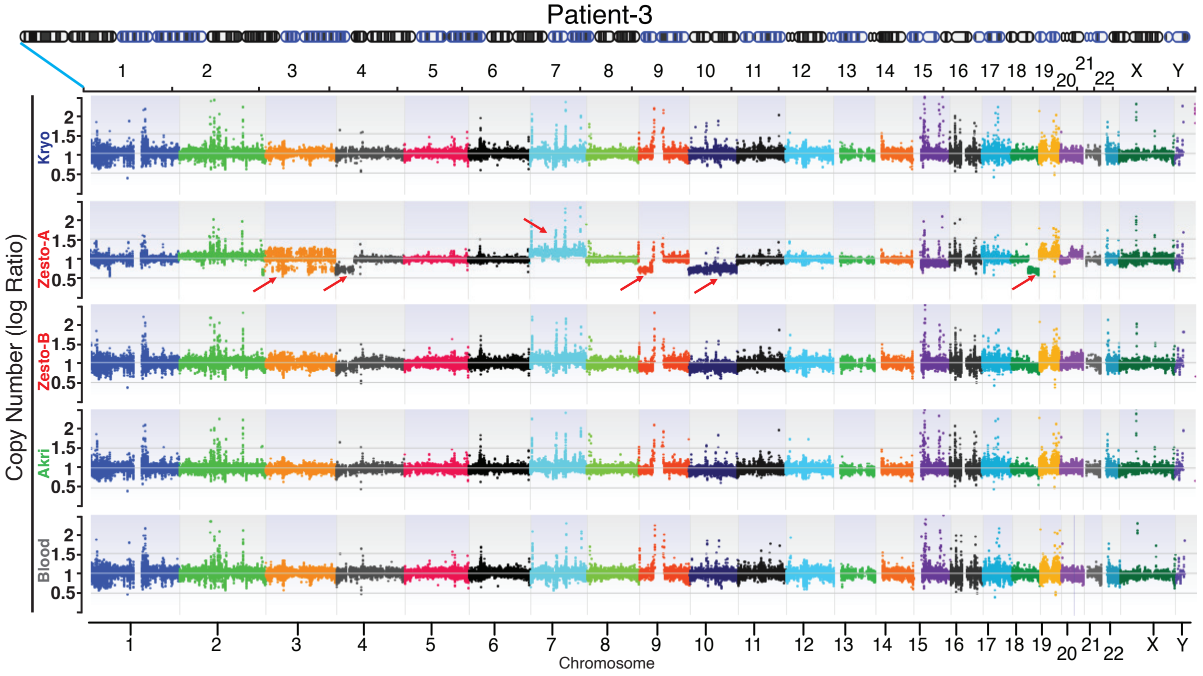


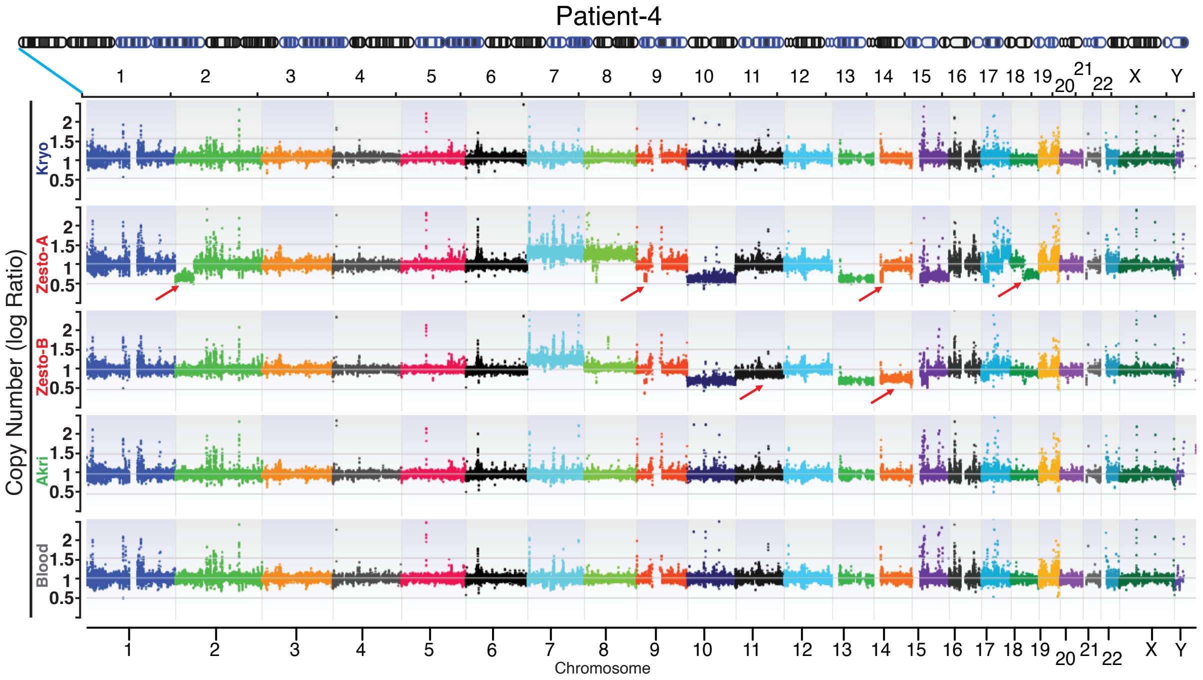


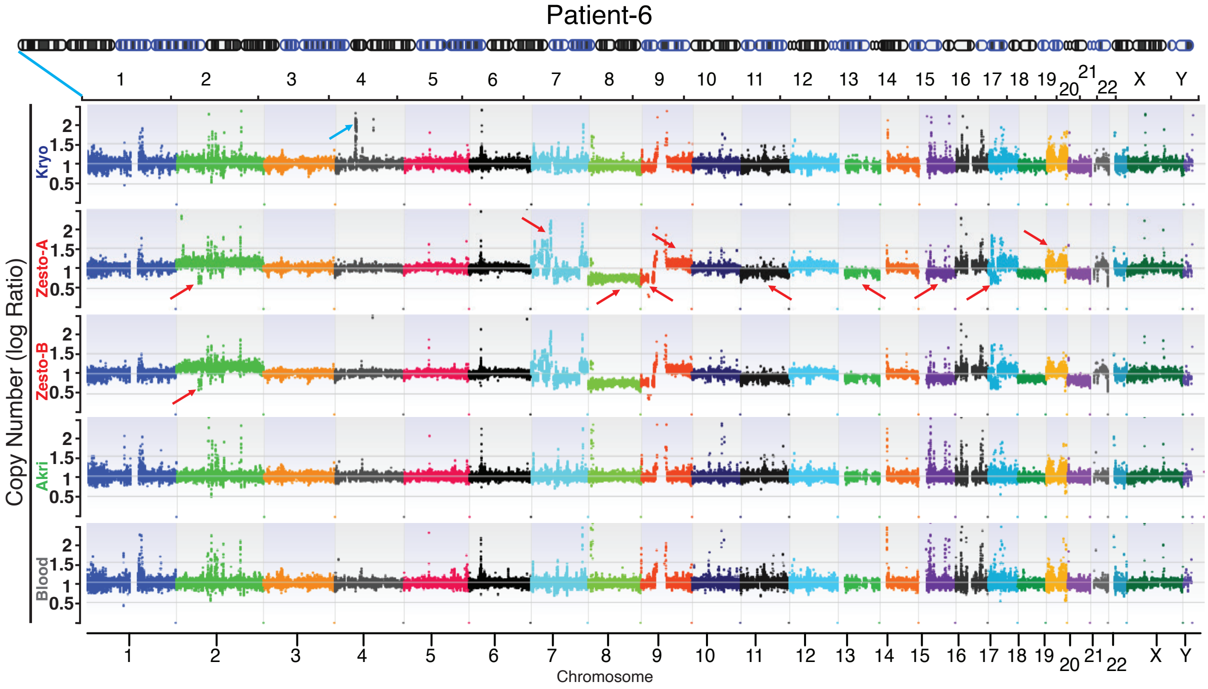

Supplementary figure S9

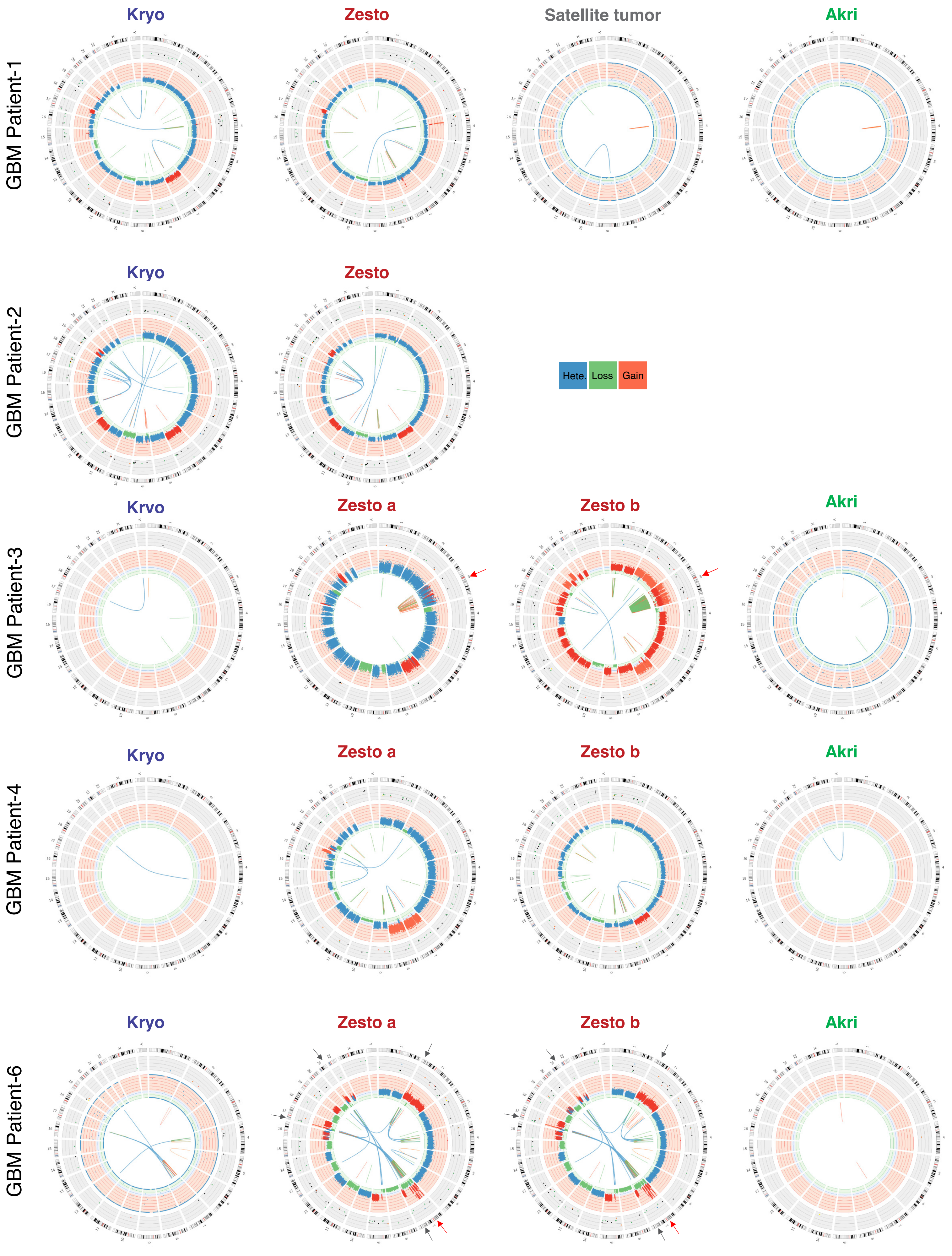

Supplementary figure S10

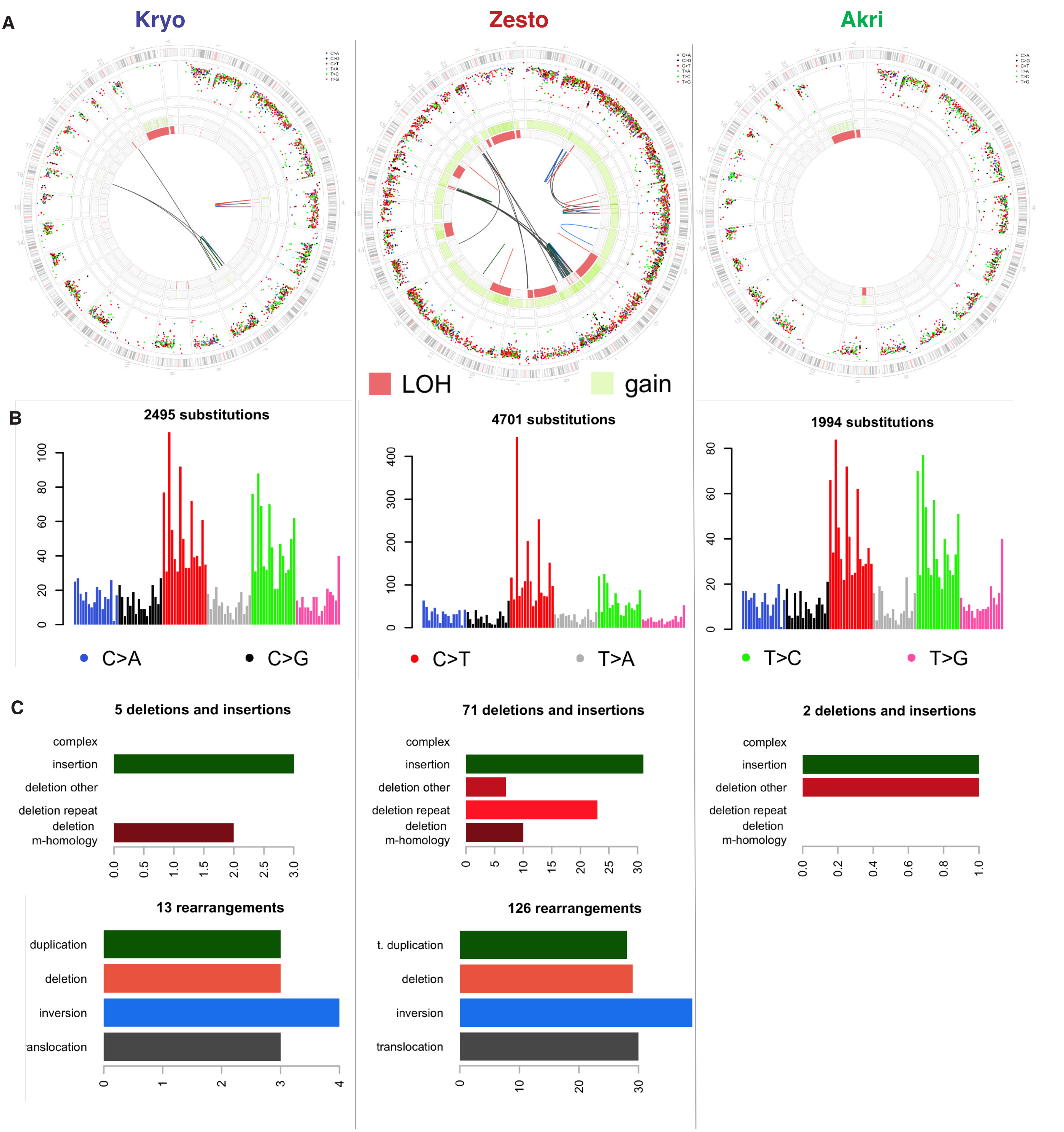

Supplementary Fig. S11

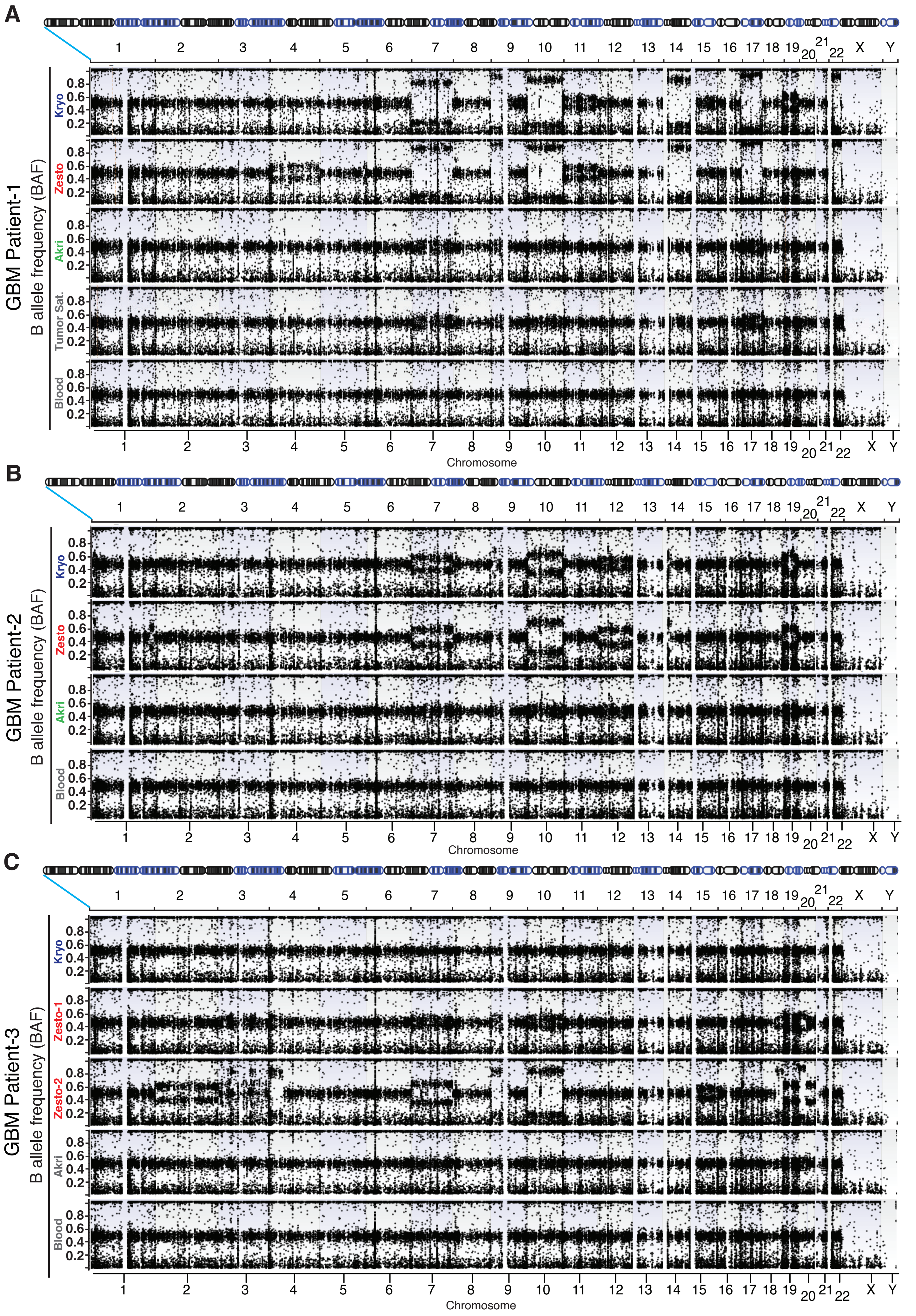

Supplementary Fig. S12

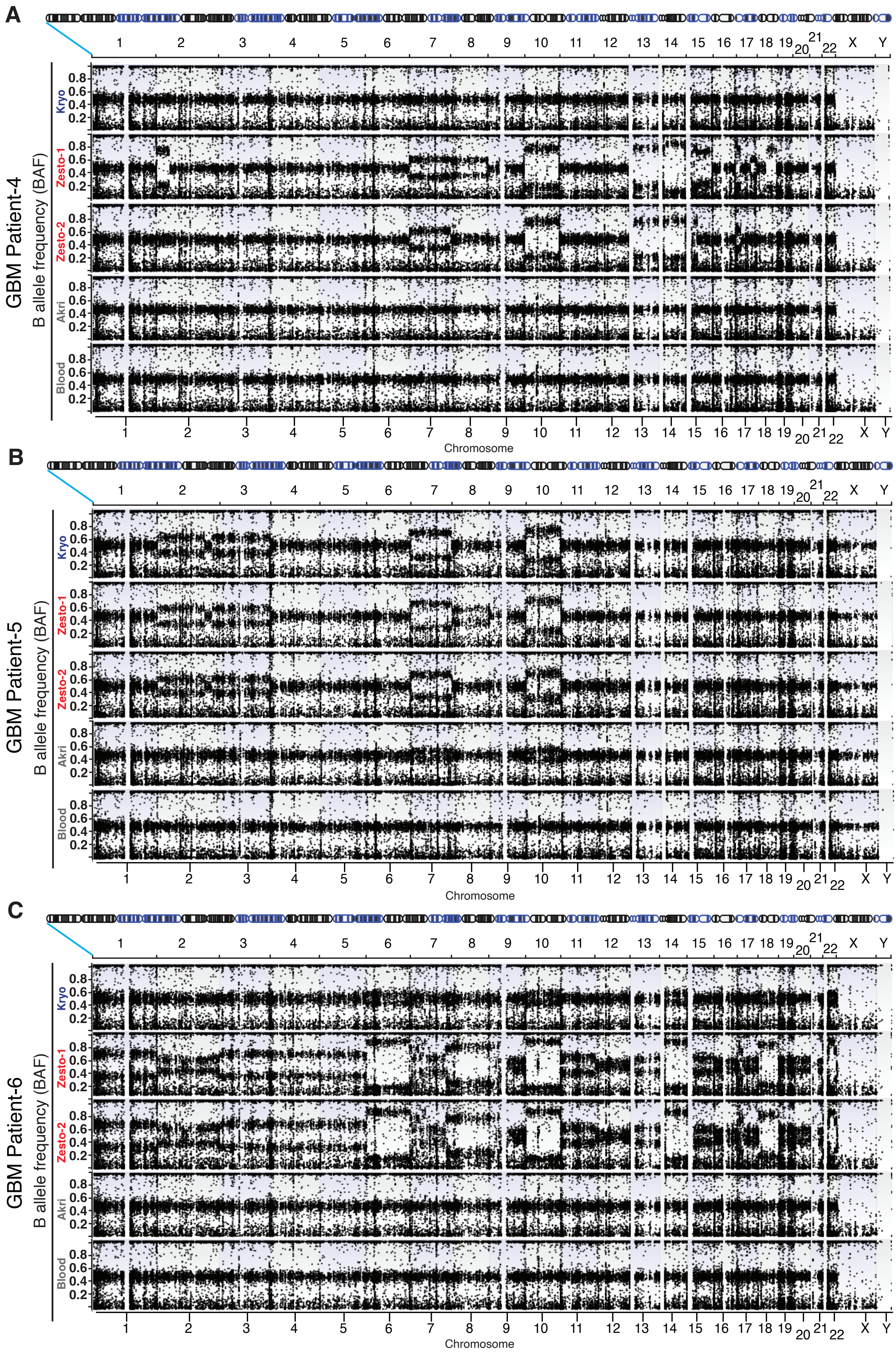
